# A simple circuit to sustain intact tumor microenvironments for complex drug interrogations

**DOI:** 10.1101/2025.10.26.684624

**Authors:** A Leila Sarvestani, Samantha Ruff, Kirsten Remmert, Martha Teke, Emily Verbus, Areeba Saif, Marcial Garmendia-Cedillos, Alex Rossi, Ashley Rainey, Krystle Nomie, Surajit Sinha, Michael M. Wach, Carrie Ryan, Stephanie N. Gregory, Priyanka Desai, Abir K. Panda, Emily C. Smith, Reed Ayabe, James D. McDonald, Shreya Gupta, Tracey Pu, Tahsin Khan, Jacob Lambdin, Kenneth Luberice, Stephanie C. Lux, Hanna Hong, Imani Alexander, Sarfraz R. Akmal, Shahyan Rehman, Sophia Xiao, Jason Ho, Justine Burke, Cathleen E. Hannah, Alexandra Livanova, Andrey Tyshevich, Stanislav Kurpe, Andrey Kravets, Basim Salem, Daniil Vibe, Arina Tkachukv, Anna Belozerova, Aleksandr Sarachakov, Evgeny Barykin, Vladimir Kushnarev, Ekaterina Postovalova, Alexander Bagaev, Noemi Kedei, Maria Hernandez, Baktiar Karim, Donna Butcher, Jennifer Matta, Michael Kelly, Jatinder Singh, Todd D. Prickett, James Hodge, Thorkell Andresson, Michael B. Yaffe, Ethan M. Shevach, David Kleiner, Jeremy L. Davis, Andrew M. Blakely, Jonathan M. Hernandez

## Abstract

Deep learning and large language models can integrate complex datasets to uncover biological insights that are often undetectable through conventional analyses. With application to translational cancer research, these computational tools have positioned 3D patient-derived tumor avatars front and center as crucial data input sources. However, a major challenge remains: the lack of standardization in media composition in 3D patient-derived tumor models unpredictably affects cell behavior and limit the utility beyond predicting treatment responses. To address this unmet need, we developed a simple, reproducible perfusion circuit system to approximate *in vivo* physiology using autologous patient plasma. With peritoneal metastases and core needle biopsies across multiple tumor histologies, we demonstrate preservation of the tumor microenvironment for up to 48 hours using multi-modal interrogation techniques. With proof-of-concept experiments, we display the system’s ability to unveil complex drug-dependent biology within this time window. Standardizable, physiologically relevant platforms for 3D patient-derived tumor avatars will yield unprecedented insights through the integration of data from broad groups of patients and the use of an expanding armamentarium of artificial intelligence capabilities.

## INTRODUCTION

Advances in computational science are re-shaping the use of human tumors in translational medicine. Deep learning and large language models can interpret and integrate the complex datasets generated by multi-omics technologies, including genomics, transcriptomics (bulk, single cell and spatial), and proteomics.^1^ These technologies offer a multilayered view of cancer with the potential to uncover critical insights into drug-dependent biology that are not currently achievable through conventional analyses. For example, while checkpoint inhibitors have been administered to tens of thousands of patients to date, our ability to understand and predict treatment response remains surprisingly limited with standard biomarker assessments (e.g. PD1/PDL-1).^2^ However, Artificial Intelligence (AI) models excel at integrating multimodal data to identify non-linear patterns and reveal hidden biology.^3^

3D patient-derived tumor avatars (3D-PTAs), such as organoids and tumor slices, are emerging as powerful tools for modeling individual human tumors. 3D-PTAs offer significant advantages over traditional models like 2D cultures or patient-derived xenografts (PDXs), including faster turnaround times, greater scalability, and compatibility with high-throughput drug screening.^4^ Most importantly, it is possible for these avatars to preserve the native tumor microenvironment (TME), allowing for dynamic, physiologically relevant testing of therapeutic responses.^5^ As an example, perfusion culture systems in pancreatic ductal adenocarcinoma can maintain both immune and stromal components, enabling evaluation of patient-specific immune landscapes and treatment effects.^6^ Organoids are the most commonly employed 3D-PTA and are currently unmatched in terms of high-throughput capacity and sustainability. However, the growth and perpetuation of organoids are dependent on the use of mouse-derived extracellular matrix substitutes or fetal calf/bovine serum, which introduce undefined extrinsic variables that unpredictably influence cell populations.^7^ Media composition has been shown to impact organoid growth rate, morphology, and sensitivity to chemotherapy or targeted agents.^8,9^ While tumor slices better preserve the complexity of the TME, the processing and perfusion can be complex and, like organoids, they require media composition that unpredictably influences the TME. Taken together, the integration of 3D-PTA readouts across various data input sources for AI analyses will be extremely challenging without standardization, including universal media preparations.

When integrated with molecular profiling (multi-omics), 3D-PTAs stand to deliver on the goal of precision medicine: matching patients to effective treatments while avoiding ineffective therapies that will only impart toxicity. However, it is crucial to realize that drugs demonstrating clinical efficacy rarely work to the satisfaction of the patients receiving them. Complete tumor regression is an extremely infrequent event in solid tumor oncology, especially for patients with metastatic disease. Far more commonly, efficacious first-line drug combinations yield partial responses (shrinkage of the tumor on advanced imaging by ≥30%) and are continued until progression, typically 6-12 months. We posit that human tumors can *also* be utilized to revise and improve drug combination regimens through detailed AI analyses. To achieve this endpoint, sustaining human tumors *ex vivo* must be <u>simple</u> to allow for broad adoption and elimination of extrinsic and unpredictable variables, <u>durable</u> to observe drug-dependent alterations, and <u>standardizable</u> to leverage the evolving advances in computational science. Herein, we crafted a perfusion circuit, termed the SMART (<u>S</u>ustained <u>M</u>icroenvironment for <u>A</u>nalysis of <u>R</u>esected <u>T</u>issue) system, to approximate normal human physiology with autologous patient plasma as a readily accessible, reproducible media preparation. We used human metastases to evaluate the circuit given that these tumors do not interfere with the intricacies of primary tumor staging, multiple tumors are often present to help account for intertumoral heterogeneity, and tumor quantity is sufficient for both research and clinical applications. Through detailed interrogation, we demonstrate that 3D TME architecture, including cell-cell interactions, were maintained to allow for detailed drug response evaluations. These advances bring us closer to realizing the full potential of patient-derived tumors as platforms for therapeutic innovation.

## RESULTS

### SMART System Development

To interrogate <u>intact</u> human metastases outside of the patient, we crafted a perfusion circuit to allow for gas/waste/nutrient exchange with tissue fixed on user-modifiable platforms that fit lock-in-key on a suspension lid **(Figure 1A-C)**. The circuit consists of inflow tubing with a rollator pump that propels plasma through a silicone membrane gas exchanger before entering a 3D-printed circular dish housing the platforms. Given the deleterious effects of shear stress on tissue viability,^10^ we determined optimal platform orientation and plasma flow rate using an iterative angle change experiment (delta 0.3 degrees for values between 0 and 45 degrees) with flow rates between 1 ml/min and 30 ml/min. (**Figure 1D, Figure S1A**). To visualize these parameters, we used a high-speed camera with light-diffracting beads added dropwise to the inflow plasma circulating at 15.5 ml/min **(Figure S1B**). Next, perfusate parameters were evaluated over time in a standard 37°C incubator. An obvious limitation of the system is the lack of red blood cells for oxygen delivery, which we sought to mitigate with the use of 100% O_2_. Pure O_2,_ however, resulted in significant basic pH derangements. To maintain a physiologic pH, a gas mixer was employed to add CO_2_ at a starting mix of 93% O_2_ and 7% CO_2_, with subsequent adjustments made according to point-of-care measurements **(Figure 1E)**. Importantly, dissolved oxygen content was not appreciably altered with the addition of CO_2_ **(Figure 1F)**. Next, sodium concentration was used to evaluate ion fluctuations over time given that derangements above and below physiologic parameters have been reported to affect immune populations.^11^ Hypernatremia secondary to evaporative water loss was evident and was corrected with continuous water infusion (starting rate of 0.5 ml/hr) using a syringe pump, with adjustments made according to point-of-care measurements **(Figure 1G)**. Finally, to minimize the risk of bacterial or fungal contamination, perfusate sampling and interventions were conducted through two in-line, 4-way stopcocks.

**Figure 1.**
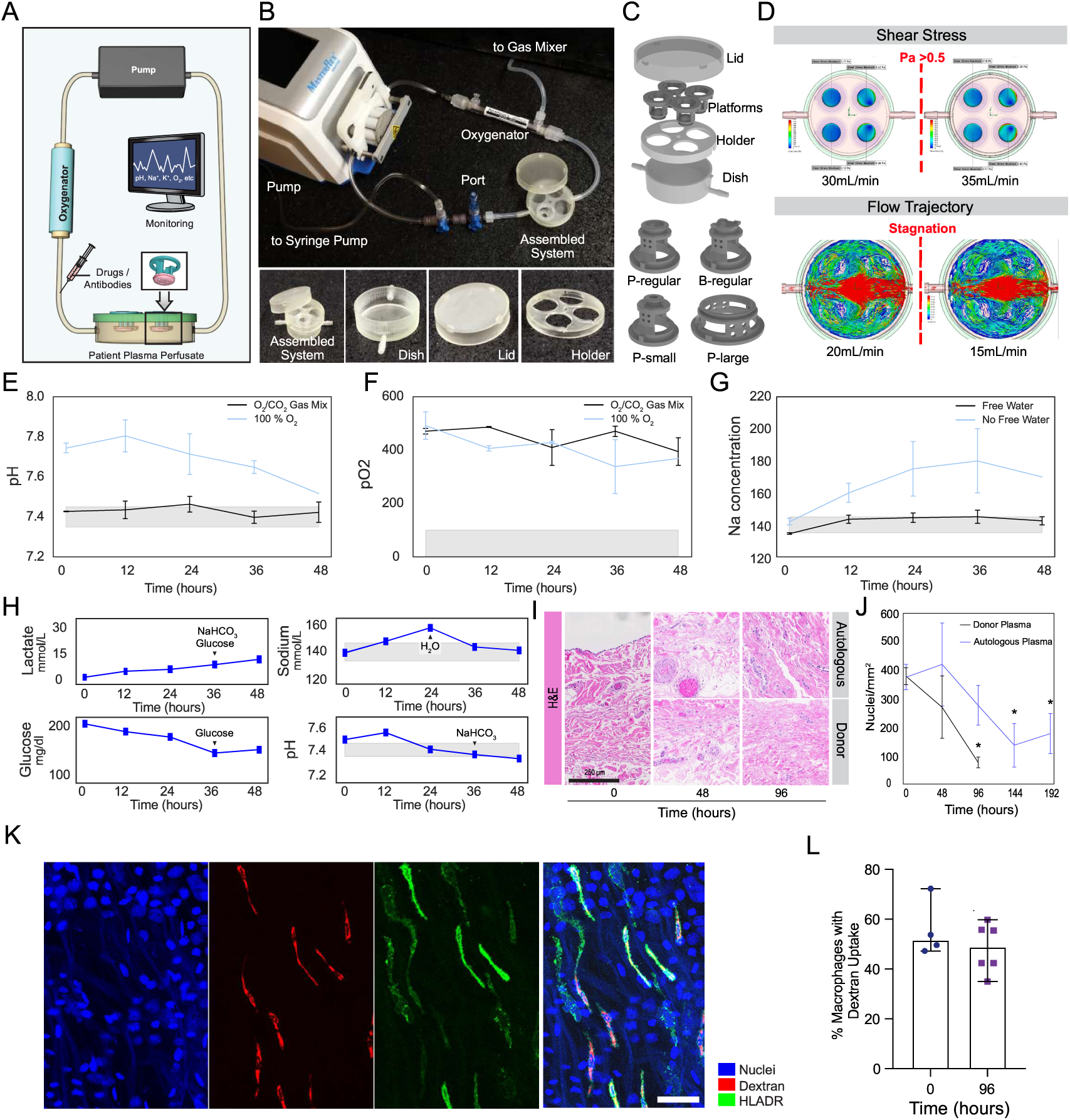
Development and evaluation of the SMART system. **A:** Schematic of the simple circuit used in the SMART system. **B:** Photograph of the SMART system circuit. **C:** Schematic of user-modifiable platforms that fit lock-in-key on a suspension lid. **D:** Determination of shear stress and flow trajectories in SMART culture dish at different flow rates. See **Figure S1A,B**. **E-G:** Graphs showing pH, dissolved oxygen (pO_2_), and sodium (Na) in autologous plasma perfusate over 48 hours in the SMART system, respectively. **H:** Graphs from a representative run showing lactate, sodium, glucose, and pH after the introduction of patient tissue into the SMART system. Arrows indicate interventions made based upon point-of-care measurements. **I:** H&E images of tissue in the SMART system perfused with autologous plasma or cross-matched donor plasma over 96 hrs. **J:** Graph of nuclei/mm^2^ measured over 192 hrs in the SMART system perfused with autologous plasma or cross-matched donor plasma. **K:** HLADR-positive macrophages (green) maintain the ability to constitutively engulf fluorescently conjugated Dextran (red). Representative image. Scale bar: 50 um. **L:** Percentage of Dextran- and HLADR-positive cells observed at 0 and 96 hrs of perfusion.

Satisfied with the control of perfusate within the measured physiologic parameters, the system’s effects on tissue were assessed. To begin, parietal peritoneum from patients undergoing risk-reducing operations (patients without current cancer diagnosis) was utilized with the rationale of minimal heterogeneity between humans for ease of repetitive testing. Of note, the tissue was acquired at the beginning of the surgical procedure to minimize the short-lived inflammatory influx that increases with operative duration.^12^ With the introduction of tissue, glucose levels decreased and lactate levels increased over time, consistent with carbohydrate metabolism. These observations led us to institute glucose and sodium bicarbonate additions (based on point-of-care measurements) and a perfusate change every 48 hrs given concerns of metabolic waste buildup without dialysis in the circuit **(Figure 1H, Figure S1C)**. The system as outlined does not allow for dynamic influx of new cell populations, and therefore, cell numbers were documented over time to select optimal experimental timepoints. With maintenance of perfusate within physiologic parameters, nuclei were evaluated every 48 hrs for a duration of 192 hrs using hematoxylin and eosin (H&E) stains. As compared to baseline, a statistically significant decrease in cell nuclei was seen at 144 hrs **(Figure 1I,J**), indicating 48-hr and 96-hr timepoints may be optimal. Importantly, short-term cultures (48–72 hrs) are sufficient to determine the immunomodulation of the tumor immune microenvironment with immunotherapy.^13^ Given the blood requirements with autologous plasma for multiple systems (e.g. control, condition A, condition B), we evaluated the use of cross-matched donor plasma as a potential substitute/supplement. However, ABO(H) histo blood group antigens are expressed on most epithelial and endothelial cells, and even “compatible” donor plasma can carry alloantibodies that activate complement on tissue surfaces. In line with expectations, perfusate composed of cross-matched donor plasma led to substantial reductions in nuclei numbers **(Figure 1I,J, Figure S1D)**, and therefore, only autologous plasma was utilized. Finally, to evaluate resident cell function over 96 hrs, we focused on macrophages, the tissue sentinels contributing to tissue homeostasis, remodeling and repair. Macrophages constantly sample their environment through actin- and phosphatidylinositol 3-kinase (PI3K)-dependent membrane ruffling, leading to the generation of internal organelles filled with extracellular fluid. This constitutive process leads to the internalization of a cell surface equivalent every 33 mins. Importantly, macrophage function is sensitive to external stimuli including hypoxia, oxidative stress, and nutrient depletion.^14,15^ To assess macrophage function over time, constitutive uptake of the metabolic substrate dextran was monitored. Accumulation of fluorescently conjugated dextran in HLADR-positive macrophages was not significantly altered over 96 hours, indicating that the system does not impact constitutive pinocytosis of metabolic substrates **(Figure 1K,L)**.

### Effect of the SMART System on TME Populations

Following physiological optimization of perfusion parameters and evaluation of normal tissue, we sought to evaluate the system’s effects on cell types present in the TME using multi-omic interrogation (**Figure S2**). To this end, we utilized complementary approaches on multiple levels: morphologically with H&E staining, transcriptionally with bulk RNA-based gene expression analysis and cell deconvolution, and at the protein level with immunofluorescence. With maintenance of physiologic parameters, metastases were evaluated from 11 patients using QuPath on H&E-stained sections (**Figure S3B-D, representative images**). As shown in **Figure 2A**, the various cell densities and levels of necrosis remained primarily unchanged, with no significant differences observed over time. Importantly, evaluation of immune cell density demonstrated few differences over the time course, indicating the effects of perfusion do not dramatically alter the density of the TME cellular components. To further confirm this finding, RNA-based cell deconvolution using the Kassandra algorithm^16^ was performed on a subset of tumors from 10 patients at baseline and after 48 hrs or 96 hrs of perfusion. The expected percentages of TME cell types across these tumor types were found to have insignificant shifts in percentages over time as demonstrated in a heatmap (**Figure 2B**). Again focusing on macrophages given their sensitivity to environmental perturbations, the predicted macrophage cell percentages primarily remained unchanged in three patients following 96 hrs of perfusion as compared to baseline: 5% vs 15 % (colon cancer), 6% vs 9.5% (mesothelioma), and 4% vs 5% (gastric cancer) (**Figure S3E**). Notably, the only cell population to shift over time was the neutrophil population, which may be expected given the reported short half-life for these cells.^17^

**Figure 2.**
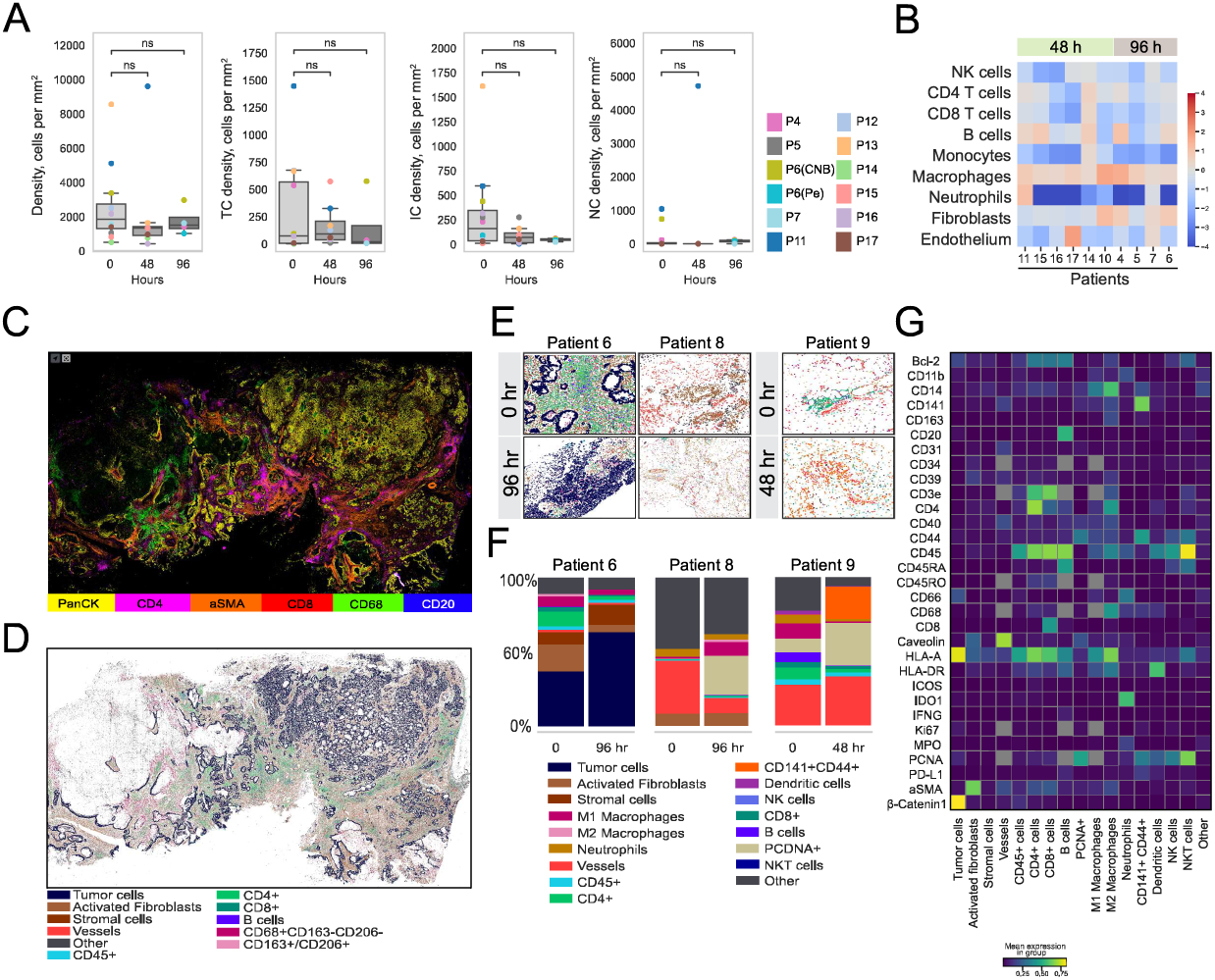
TME preservation in the SMART system. **A:** Boxplots depicting overall cell density, tumor cell (TC) density, immune cell (IC) density, and necrotic cell (NC) density of tumors perfused in the SMART system for 48 or 96 hrs. In the boxplots, the bottom whisker indicates the maximum value or 75th percentile + 1.5 interquartile range (IQR); the top whisker indicates the minimum value or 25th percentile − 1.5 IQR. Peritoneum (Pe) and core needle biopsy (CNB). See Figure S3B-D. **B:** Heatmap of relative percentages of cell populations derived from RNA-based cell deconvolution at 0, 48 and 96 hrs. **C-D:** Representative multiplex immunofluorescence image and cell segmentation image of a tumor from Patient 6 at 0 hr. See **Figure S3E,F**. **E:** Representative multiplex immunofluorescence and cell typing images of tumors from Patient 6, Patient 8, and Patient 9 at the indicated timepoints. **F:** Bar graphs depicting cell percentages derived from cell typing of the patient tumors perfused in panel E. **G:** Heatmap of the cell markers across all cells showing all cell populations detected for Patient 6, Patient 8, and Patient 9 combined.

To delve deeper, we employed multiplex immunofluorescence (MxIF) to further identify cell populations using tumors from three additional patients (goblet cell appendiceal adenocarcinoma, rectal adenocarcinoma, or peritoneal mesothelioma) and an orthogonal AI-driven cell typing approach,^18,19^ which allows for TME subpopulations to be accurately quantified. We utilized tumors from the patient with rectal adenocarcinoma as proof-of-concept for the ability of this AI approach to quantify populations of different cell types. In doing so, we identified subpopulations present within the TME and demonstrated their stability during perfusion (**Figure S3 F,G**). For example, multiple tumor cell subpopulations were identified using E-cadherin and β-catenin, and over 15 minor immune populations were identified based on the detection of different combinations of markers, providing confidence in the accuracy of cell identification. **Figure 2C** depicts a representative multiplex immunofluorescence image. A machine learning (ML)-based segmentation algorithm was then employed for cell typing across the generated images for each tumor (**Figure 2D**, representative figure). As shown, various cell populations, including B cells, activated fibroblasts, stromal cells, CD4^+^ cells, and CD8^+^ cells, were identified based on marker expression detection. Overall, the cell typing after perfusion confirmed that various immune cell populations remained intact (**Figure 2E,F**). A heatmap was created based on the cell typing analysis for tumors from each of the three patients combined, depicting cell populations with various markers (**Figure 2G**).

### Effect of the SMART System on TME Interactions

#### Architecture

The persistence of TME cell populations over 96 hrs of perfusion does not necessarily indicate preserved cellular interactions and contacts between the cell populations. We therefore employed AI-based analysis of MxIF imaging to identify cellular neighborhoods and communities and assess the stability of this architecture over time. Using tumors from six patients, cellular neighborhoods and communities were categorized into nine major zones (**Figure 3A, Figure S4A-D**). As shown in the analyzed sections, the vast majority of the identified communities persisted throughout perfusion, with a few new communities identified. A median percent change ranging from 2-8% in community proportions was observed over time (**Figure 3B**). In addition, permutation testing was employed by analyzing the non-randomness of the cellular contacts.^20^ While some cell-cell contacts changed from baseline after 96 hrs of perfusion, including the identification of new contacts and the loss of previous contacts, most cellular interactions were sustained over time (**Figure 3C)**. For example, significant contacts were maintained after 96 hrs among CD4^+^ T cells, CD8^+^ T cells, and T regulatory cells in tumors from a patient with peritoneal mesothelioma. In another example, CD4^+^ T cells and macrophages remained in contact after 96 hrs of perfusion in tumors from a patient with appendiceal adenocarcinoma. Across all samples, 77.4% of the cellular contacts present at baseline persisted following 96 hrs of perfusion. Next, the non-cellular TME was assessed using Masson’s Trichrome stain in Patient 6 and Patient 7, revealing no significant differences in optical density as a result of perfusion (**Figure S4E**). In summary, the spatial architecture results indicate that the TME remains largely intact, albeit not entirely unchanged, following 48 hrs or 96 hrs of perfusion.

**Figure 3.**
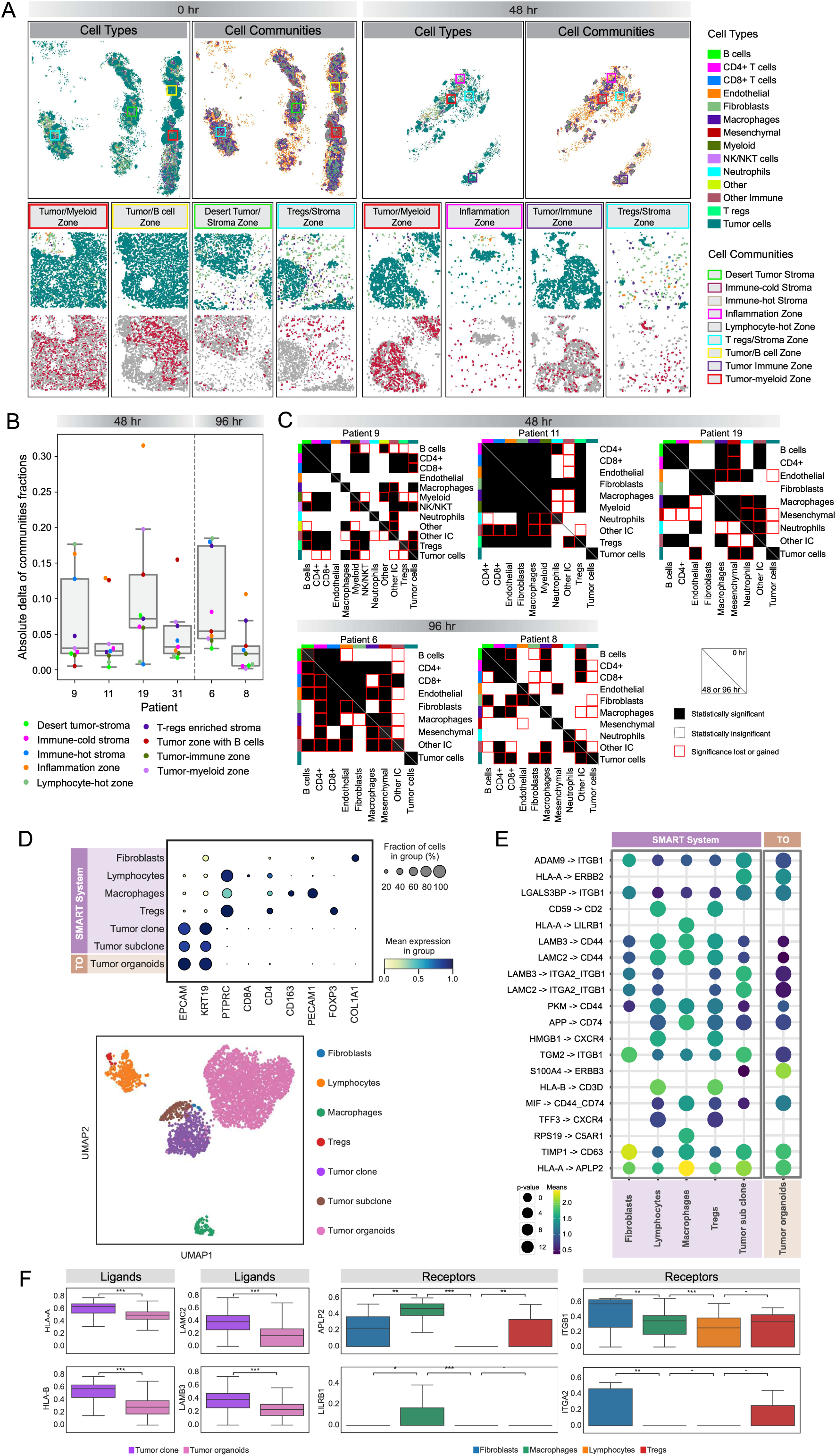
TME neighborhood and community analysis in the SMART system. **A:** Cell typing and community spatial plots for tumors from Patient 11 (full tissue-top; tissue regions-bottom) at 0 hr and after 48 hr of perfusion. See **Figure S4A-D**. **B:** Box plots of absolute delta between communities observed at 0 hr and after 48 hr (Patient 9, Patient 11, Patient 19, and 31) or 96 hr (Patient 6 and Patient 8) of perfusion. In the boxplots, the bottom whisker indicates the maximum value or 75th percentile + 1.5 interquartile range (IQR); the top whisker indicates the minimum value or 25th percentile - 1.5 IQR. **C:** Heatmaps showing statistically significant (non-random) cellular contacts after 48 hr (Patient 9, Patient 11, and Patient 19) or 96 hr (patient 6 and Patient 8) of perfusion. **D:** Top-Dot plot with the size of the circles representing the percentage of the listed cell types present in the SMART system or tumor organoid and the mean expression levels of the listed genes shown in blue color scale. Bottom-UMAP plot of the cell types identified in the SMART system and tumor organoid. See **Figure S5B**. **E:** Dot plot with the size of the circles representing the p-value significance of a potential ligand-receptor interaction and the color scale representing the expression level of both the ligand and receptor in each cell type. **F:** Boxplots depicting expression levels of the ligands and receptors in the delineated cell types. In the boxplots, the bottom whisker indicates the maximum value or 75th percentile + 1.5 interquartile range (IQR); the top whisker indicates the minimum value or 25th percentile - 1.5 IQR.

#### Ligand Receptor Interactions

To frame the importance of TME architecture on the tumor cells present, we derived organoids from a patient whose tumor was also perfused in the system for 96 hrs. Organoids are a convenient model, but the tissue constructs generally lack the non-tumor cell components of the TME unless special preparations are undertaken. We interrogated tumors (following 96 hrs of perfusion) and organoids derived from a patient with rectal adenocarcinoma using multiple molecular and imaging approaches such as scRNA-seq, bulk RNA-seq, and multiplex immunofluorescence. The organoid model was created utilizing previously established protocols,^21,22^ and after 4 passages, was interrogated similarly with tumor perfused in the system. scRNA-seq showed, not surprisingly, that the organoid was only comprised of tumor cells whereas tumor perfused in the system consisted of TME cells such as T regulatory cells, macrophages, lymphocytes, fibroblasts (**Figure 3D**). The scRNA-seq analyses also showed two major epithelial cell populations, denoted clone and subclone (**Figure S4F**), which differed transcriptionally from tumor cells in the organoid population (**Figure S4G**).

Ligand-receptor interactions are critical to the intricate interplay between tumor cells and the immune microenvironment and can be inferred with scRNA-seq. Using CellPhoneDB,^23^ ligand-receptor interactions were predicted based on the expression of specific ligands and receptors across the cell types identified in perfused tumors **(Figure 3E)**. Notably, while the organoid itself was comprised only of tumor cells, we utilized TME cell populations from the SMART sample to predict potential cell–cell interactions. This allowed us to assess what possible interactions might occur between organoid tumor cells and TME components, rather than restricting the analysis to tumor–tumor interactions alone. The ligand-receptor interactions found in the perfused tumor were numerous, with key interactions necessary for an immune response. For example, TIMP1-CD63 and HLA-A-APLP2 interactions were strongly predicted to occur between the tumor and fibroblasts and macrophages, respectively. Importantly, tumor cells following perfusion showed multiple predicted interactions with the major cell populations within the system (**Figure 3F)**. SMART tumor cells had higher expression of ligands of Tumor-TME interactions than the organoid, highlighting the importance of an intact TME to maintain the tumor cell transcriptome.^24^ These predicted interactions in conjunction with architecture data suggest that the TME and its complex intercellular interactions are present during perfusion in the system.

### Effect of the SMART System on Proteo-transcriptomics and Cell Function

#### Proteo-transcriptomics

Neutrophil death can trigger inflammatory responses and/or activate stress response pathways that may influence intra- and inter-cellular signaling and cell phenotypes. Given neutrophil loss coupled with the observed alterations in TME architecture during perfusion, we sought to further interrogate the impact of the system on the TME over time with increasingly sensitive metrics.^25^ First, we utilized bulk RNA-seq to conduct gene expression analysis across tumors from three patients. Pathway analyses did not demonstrate significant changes over 96 hrs, indicating that perfusion does not greatly induce the activation of stress response genes detectable at the mRNA level (**Figure 4A**). Moreover, differential gene expression across tumors from these three patients showed differences in only a minimal number of genes (450 genes out of ∼16,000, 2.8%) after perfusion (**Figure 4B**). Next, we performed proteomic analyses via mass spectrometry, as proteins are degraded in tightly regulated dynamic processes that play critical roles in maintaining cellular homeostasis and response to environmental cues. We evaluated tumors from the patient with gastric cancer (after 96 hrs of perfusion) who was also included in the transcriptomic analysis. Interestingly, these tumors demonstrated more proteomic changes than anticipated based upon the transcriptomic data (**Figure 4C, top**). To evaluate further, we analyzed tumors from another patient (ovarian cancer) after 48 hrs of perfusion. As shown in the volcano plot (**Figure 4C, bottom),** only a small portion of this analyzed proteome was significantly altered during 48 hrs of perfusion. Pathway analysis was performed on data from both patients revealing marked upregulation of coagulation and complement pathways, which may be expected given that static blood remains in the capillaries/microvasculature during perfusion (**Figure 4D**). Interestingly, consistencies across both time points included upregulation of EMT, KRAS signaling, and angiogenesis, whereas consistent downregulation involved fatty acid metabolism, oxidative phosphorylation, heme metabolism, and adipogenesis. Although not transcriptionally detected, the lack of an oxygen carrier could have contributed to the increase in angiogenesis proteins and similarly could have contributed to decreased oxidative phosphorylation. These processes, if present, are likely to affect the TME globally. Fluctuations in oncogenic signaling, however, may alter the balance of tumor immune cross-talk in unpredictable ways, which may be problematic. For example, Kras-driven signaling suppresses MHC-I dependent antigen presentation and induces expression of PD-L1, while decreased PI3K/AKT/mTOR pathway output can deplete T regulatory cells and enhance cytotoxic T cell effector functions.^26–28^

**Figure 4.**
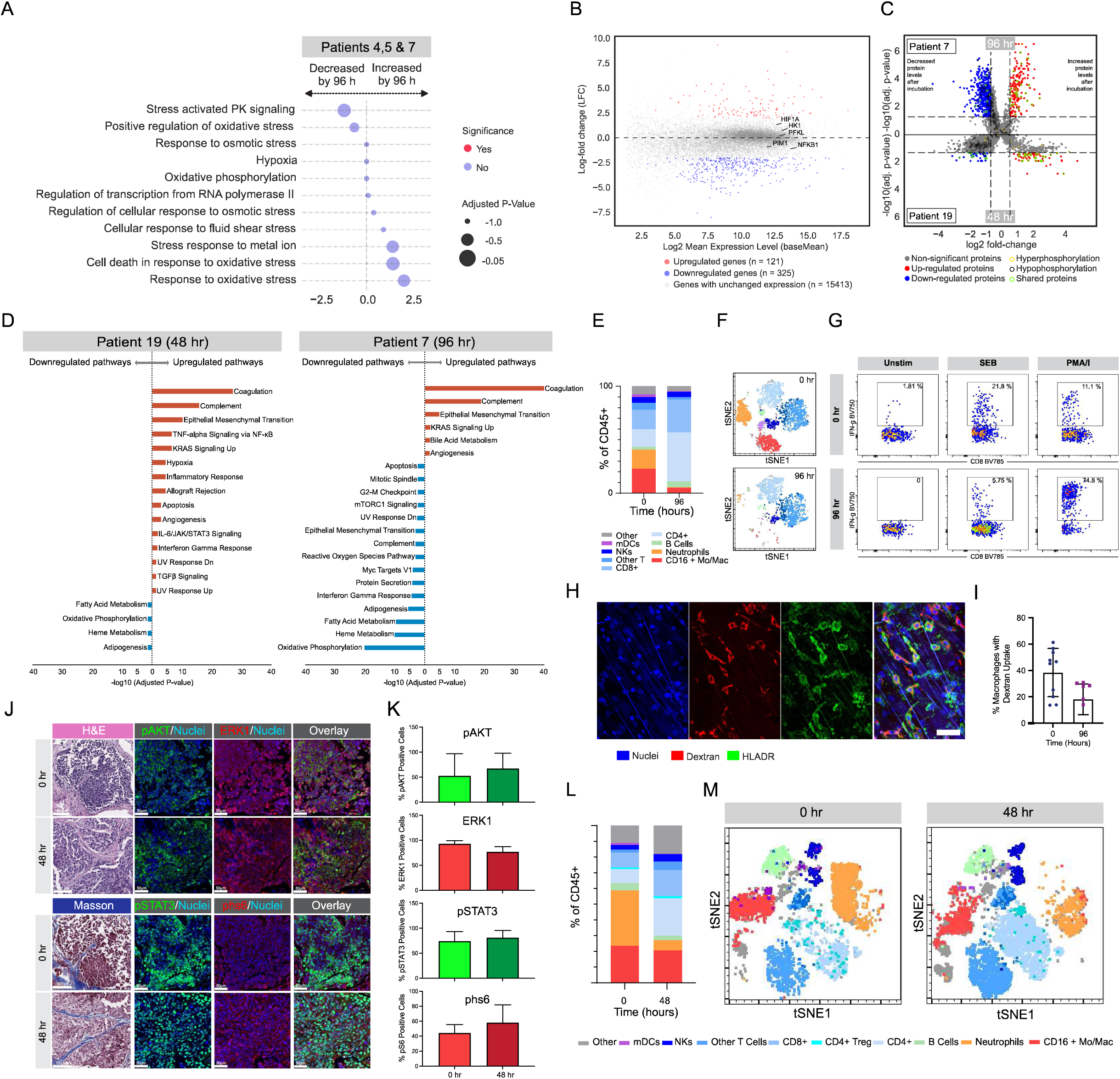
Functional analysis of SMART system tumors. **A:** Dot plot showing significance of mRNA expression changes related to the listed pathways for tumors from Patient 4, Patient 5, and Patient 7 after 96 hrs of perfusion. **B:** Scatter plots depicting changes in gene expression for tumors from Patient 4, Patient 5, and Patient 7 after 96 hrs of perfusion. **C:** Scatter plots showing change in protein expression for tumors from Patient 7 (top) and Patient 19 (bottom) after 96 hrs or 48 hrs of perfusion, respectively. **D:** Pathway analysis of protein data in Panel **C**. **E:** Immune populations were identified in unstimulated wells and quantified as % of CD45^+^ events after 96 hr of perfusion. **F:** Unsupervised visualization of immune subsets by t-SNE. **G:** CD8^+^ T cell responsiveness was assessed at baseline (0 h, top) and following perfusion (96 hr, bottom) with either SEB, PMA/I, or were unstimulated for 3 hrs in the presence of Brefeldin A and were evaluated for IFNγ production. See Figure S5C. **H:** HLADR-positive macrophages (green) within peritoneal tumor biopsies maintain ability to constitutively engulf fluorescently conjugated Dextran (red) in the perfusion circuit over 96 hr. Representative image. Scale bar: 50 um. **I:** Percentage of Dextran- and HLADR-positive cells observed at 0 hr and 96 hr. **J:** Assessment of the phosphorylation status of AKT, ERK1, STAT3 and RPS6 in tumor samples post resection and following 48 hr in the perfusion circuit by immunofluorescence (blue: nuclei, green: pAKT and pSTAT3, red: pERK1 and pRPS6); H&E or Masson’s Trichrome Stain included. **K:** Quantitative HALO analysis of sample sets depicted in **(J)** shows no significant decrease or increase in the phosphorylation status of AKT, ERK1, STAT3 and RPS6 following 48 hr of perfusion. **L:** Immune subsets present following 48 hr of perfusion by flow cytometry. Gated composition of immune subsets. Immune populations are quantified as % of CD45^+^ events. **M:** Unsupervised visualization of immune subsets by t-SNE following 48 hr of perfusion.

#### TME cell functional assessment

Given our concerns about greater alterations in protein signaling after 96 hrs of perfusion coupled with the documented loss of neutrophils and some loss of cell-cell contacts, we decided to re-examine population viability over time using flow cytometry, which inherently requires dissociation of the tumor. Relative to tumor dissociated at baseline, tumor dissociated after 96 hrs of perfusion demonstrated depletion of neutrophils (consistent with transcriptomic-based cell deconvolution, see **Figure 2B**) and monocyte/macrophage subsets (**Figure 4E,F**). Although macrophage loss with dissociation has been reported previously,^29^ these results indicate increased fragility as a result of perfusion in the system. Of note, monocyte/macrophage depletion was not observed with pathology sections from prior samples **(Figure 2E-G)**, highlighting the importance of metrics beyond “roll call” to determine culture conditions on cell populations. Along this line, we investigated the ability of TME T cells to respond to stimulus. We again employed flow cytometry after 96 hours of perfusion in the system to demonstrate that T cells retained the ability to respond to PMA/Ionomycin and through the T cell receptor following stimulation with staphylococcal enterotoxin B (SEB; CD8^+^ T cells **Figure 4G**, CD4^+^ T cells **Figure S5A**). This result was largely expected even without exogenous IL-2 addition to the media, given prior *in vitr*o evaluation of T cell response to checkpoint inhibitors^30^ as well as the documented efficacy of *in vitro* expanded tumor reactive T cells to treat solid tumors.^31^ We next evaluated macrophage environmental sampling, which is a PI3K-dependent process sensitive to perturbations in external stimuli.^15^ Importantly, this evaluation identified retained but diminished (∼50%) accumulation of Dextran-positive internal organelles, again suggesting the potential for functional compromise over 96 hrs of perfusion (**Figure 4H,I**).

Given these differences between 0 and 96 hrs, we more closely evaluated tumor perfused in the system for 48 hrs. First, we assessed signaling pathways directly by employing phospho-immunofluorescence staining to gauge major oncogenic driver readouts including AKT, ERK1, RPS6, and STAT3 for tumors from the patient included in the 48-hr mass spectrometry experiment (**Figure 4C**). As compared to baseline, no differences were observed with tumor cell pAKT (52.5% vs. 67.6%, p=0.73), pERK1 (93.0% vs. 77.0.09), pRPS6 (44.31% vs. 58.1%, p=0.56), or pSTAT3 (74.29% vs. 81.1%, p=0.73) (**Figure 4J,K**). We then dissociated tumors following 48 hrs of perfusion in the system, which revealed only partial neutrophil depletion and retention of monocyte/macrophage populations **(Figure 4L,M)**. Taken together, the employed multi-omic approach strongly suggests that any impact on cell populations or their phenotypes is minimal at 48 hrs, while demonstrating the power of leveraging multi-omic technology, paving the way for drug evaluations.

### Proof-of-Concept Drug Perfusions in the SMART System

#### Anti-EGFR Therapy

A major impetus for creating a system with autologous plasma and intact TME was to preserve *in vivo* spatial relationships for interrogation of complex crosstalk between tumor and immune cells. In a proof-of-principle experiment, we perfused tumors from two patients with colorectal cancer (one with a *KRAS* wild-type tumor and the other with a *KRASG12S* mutant tumor) with the addition of an anti-EGFR monoclonal antibody (cetuximab) and analyzed post-perfusion tumors with bulk RNA-seq and Xenium single-cell-level probe based spatial transcriptomics. In *KRAS* wildtype colorectal tumors, cetuximab binds to EGFR and reduces downstream signaling, resulting in clinical improvement. In *KRAS*-activating mutant colorectal tumors, the *KRAS* mutation bypasses the EGFR signaling pathway, causing cetuximab to be ineffective and potentially harmful. Following 48 hrs of perfusion with cetuximab, the mRNA expression of MAPK, EGFR, PI3K trended toward reductions in the *KRAS-*wildtype CRC tumors, which was opposite of that observed in the *KRAS-*mutant CRC tumors **(Figure 5A**). Intriguingly, the M2 macrophage signature increased after cetuximab treatment in the *KRAS* mutant tumors but not in wild-type tumors **(Figure 5B**), consistent with prior reports of KRAS-mutant CRC tumor cells reprogramming macrophages to a TAM-like phenotype via tumor-derived CSF2 and lactate production in response to EGFR inhibitors.^32^ M2 macrophages are known to inhibit anti-tumor immune responses and promote tumor progression and importantly have been linked to cetuximab resistance in colorectal cancer.^33^

**Figure 5.**
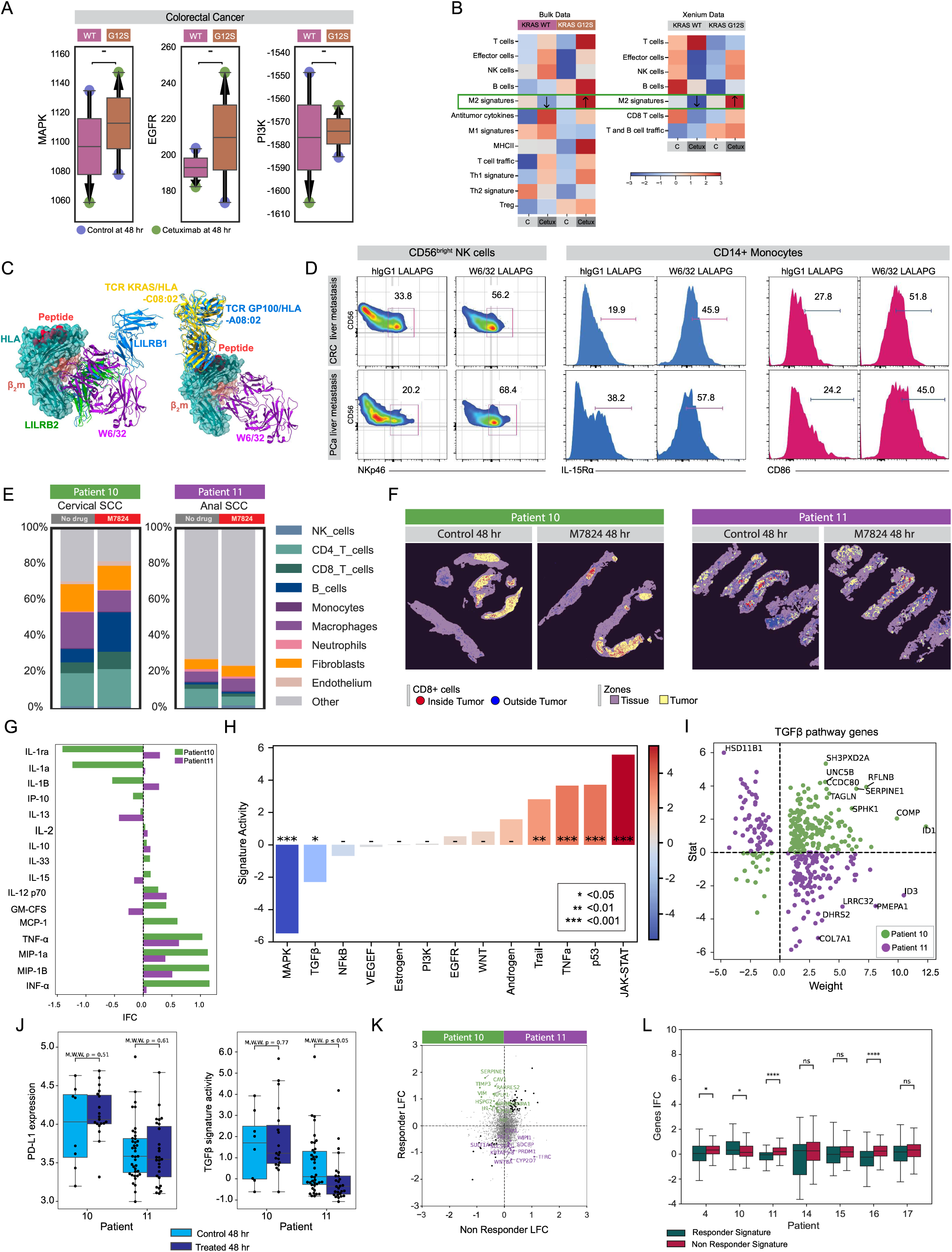
Biological effects of drug interrogation on SMART system tumors. **A:** Boxplots showing changes in mRNA expression of MAPK, EGFR, and PI3K after 48 hr of perfusion with control or cetuximab for tumors from Patient 30 and Patient 31. Arrows denote an increase (up) or decrease (down) in expression after treatment with cetuximab or control. **B:** Heatmap of mRNA expression signatures from Patient 30 and 31 tumors treated with cetuximab based on bulk RNA-seq and Xenium single cell spatial transcriptomics. **C:** Pan-anti-HLA mAb W6/32 competes for binding sites of LILRB1 & LILRB2 but not for TCR. For comparison of crystallographically defined binding footprints, the coordinates of the HLA H of W6/32/HLA-B*27:05 complex (7T0L) were superposed on the HLA H chain of the indicated structures using Chimera X (https://www.rbvi.ucsf.edu/chimerax/). Left: LILRB1/HLA-G (6AEE), LILRB2/HLA-G (2DYP). Right: TCR/HLA-A*02 (6VMA), TCR9a/HLA-C*08:02 (6ULN). For HLA molecules, only HLA-B*27:05 (teal), peptide (red) and is β_2_m (salmon) are shown. W6/32 Fab is violet; GP100 HLA-A*02 TCR is blue and KRAS HLA-C*08:02 TCR9a is yellow; LILRB1 is blue and LILRB2 is green. **D:** Flow cytometric analysis of the expression of NKp46 on NK cells, IL-15Ra and CD86 expression on CD14^+^ monocytes from TILs isolated from metastatic colorectal cancer (TOP) and metastatic pancreatic cancer (BOTTOM) perfused for 48 hr in presence of control hIgG1LALAPG or W6/32LALAPG monoclonal antibodies. **E:** Bar graph of cell percentages derived from cell deconvolution of RNA-seq performed on tumors from Patient 10 and Patient 11 treated with bintrafusp alpha or control. **F:** Representative tissue mask images based on low-plex protein analysis showing the presence of CD8^+^ cells inside and outside of the tumor zones of tumors from Patient 10 and Patient 11 treated bintrafusp alpha or control. **G:** Bar graph showing differences in cytokine levels between control-treated and bintrafusp alpha -treated tumors from Patient 10 and 11 tumors at 48 hr. **H:** Bar graph showing differences in RNA-based signature activity between control-treated and bintrafusp alpha -treated tumors from Patient 10 and 11 tumors. **I:** Scatter plot of changes in expression of TGF-β pathway-associated genes based on GeoMx analysis of bintrafusp alpha-treated versus control-treated tumors from Patient 10 and 11. **J:** Boxplots depicting PD-L1 expression and TGF-β signature activity in tumors from Patient 10 and Patient 11 tumors treated with bintrafusp alpha or control. Each dot represents detection at a unique RO I in the tissue based on GeoMx analysis. In the boxplots, the bottom whisker indicates the maximum value or 75th percentile + 1.5 interquartile range (IQ R); the top whisker indicates the minimum value or 25th percentile - 1.5 IQ R. Significance was calculated with non-adjusted Mann- W hitney- W ilcoxon non-parametric U test. **K:** Volcano plot of responder/non-responder signature based on expression changes derived from GeoMx DSP data after bintrafusp alpha treatment of tumors from Patients 10 and Patient 11. **L:** Boxplots showing expression of bintrafusp alpha Responsive (R) and non-responsive (NR) signatures applied to the combined bulk RNA-seq of tumors at baseline and after bintrafusp alpha treatment. In the boxplots, the bottom whisker indicates the maximum value or 75th percentile + 1.5 interquartile range (IQR); the top whisker indicates the minimum value or 25th percentile − 1.5 IQR. Significance was calculated with non-adjusted Mann-Whitney-Wilcoxon non-parametric U test.

#### Anti-MHC-I Therapy

With the use of the system presented herein, we recently reported the development of a humanized pan-anti-MHC-I antibody that blocks the engagement of leukocyte inhibitory like receptors (LILRs), which are expressed on all epithelial cells including tumor cells.^34^ Since MHC-I is indispensable for “self-recognition,” we hypothesized that blocking MHC-I would generate a “missing-self” response. We have concurrently been developing a second pan-anti-MHC-I chimeric antibody, W6/32LALAPG, which binds to the overlapping a3b2 site of MHC-I, thereby blocking the engagement of LILRs with MHC-I without impacting TCR binding **(Figure 5C)**. W6/32LALAPG was first incubated with PBMCs isolated from healthy donors or patients with CRC or pancreatic cancer to demonstrate markedly increased IL-15 signaling, measured CD86 expression on monocytes and pNKp46 expression on NK cells **(Figure S5B).** While NKp46 expression represents activated NK cells primed to kill tumor cells, expression of CD86 costimulatory molecule on monocytes is indispensable for T cell activation and antigen presentation along with TCR signaling.^35,36^ To evaluate the effect on tumor-infiltrating lymphocytes (TILs), we perfused tumors from these two patients (CRC and pancreatic cancer) with W6/32LALAPG for 48 hours. As compared to control treated tumors, W6/32LALAPG treated tumors demonstrated strong activation of IL-15 signaling with activation of immune markers NKp46 and CD86 **(Figure 5D)** similar to PBMCs **(Figure S5B)**.

#### Bifunctional Anti-PD-L1/TGF-β trap Therapy

Bintrafusp alpha is a first-in-class bifunctional fusion protein comprising a PD-L1 antibody combined with the extracellular domain of TGF-βRIIα β RIIα, creating a TGF-β “trap.” To evaluate its effects on the TME, we treated tumors from two patients with HPV-positive metastatic genitourinary cancers. One patient with cervical SCC (Patient 10) was a clinical responder to the drug but was forced to cease treatment after developing autoimmune adrenal insufficiency, whereas the other patient had anal SCC (Patient 11) and demonstrated a short interval of disease stabilization on the drug before rapid disease progression. Following perfusion in the SMART system with bintrafusp alpha, tumors from both patients were interrogated using multiple modalities to better understand the discrepancy in response. Of note, perfusate TGF-β levels were substantially reduced within 5 mins of drug administration, similar to that observed *in vivo* **(Figure S5C)**. Cellular deconvolution using the Kassandra algorithm applied to bulk RNA-seq demonstrated a greater number of immune cells within tumors from Patient 10, including CD8^+^ and CD4^+^ T cells, B cells, and macrophages (**Figure 5E**). This difference in immune cell composition was further confirmed using imaging that showed higher immune cell infiltration within the tumor regions and increased cytokine expression as compared to tumors from Patient 11 (**Figure 5F,G**). Interestingly, although TGF-β gene expression levels were reduced after treatment with bintrafusp alpha in the system (**Figure 5H,I**), TGF-β associated genes were not significantly different between Patient 10 and Patient 11 after treatment in the SMART system (**Figure 5J**), suggesting that the response relied heavily on the PD-L1 inhibitory component of bintrafusp alpha. The expression of the PD-L1-based network, including PD-L1, has been shown to increase after immune checkpoint blockade targeting the PD-L1/PD-1 axis in responders, suggesting an alteration in the immune response. Indeed, higher PD-L1 expression was observed in tumors from Patient 10 as compared to tumors from Patient 11 after treatment (**Figure 5J**).

Using GeoMx DSP (Digital Spatial Profiling) analysis, responder and non-responder signatures were derived based on the changes in tumors from Patient 10 and Patient 11, respectively, after bintrafusp alpha or control perfusion (**Figure 5K**). These associations were then applied to the bulk RNA-seq of tumors combined from five additional patients were at baseline and after treatment with bintrafusp alpha for 48 hrs (**Figure 5L**). Interestingly, the results show that the tumor expression profiles most closely resembled the profile of the non-responder Patient 11, suggesting that these additional patients may not have responded clinically to bintrafusp alpha. Together, these findings show that the SMART system is an extremely useful tool to interrogate the downstream effects of novel agents in an intact TME, which can help drive rational drug combinations.

## DISCUSSION

As our understanding of cancer biology advances, it has become increasingly evident that the tumor microenvironment (TME) plays a central role in tumor progression, immune evasion, and therapeutic efficacy and/or resistance.^37,38^ The integration of cytotoxic chemotherapy, immunotherapy, and targeted therapies has revealed unexpected cross-talk between oncogenic and immunologic signaling networks, further underscoring the TME’s active and dynamic role. Consequently, traditional preclinical models fail to capture this complexity. Common models, such as 2D cell culture, patient-derived xenografts, and murine models lack key components of the native TME, including human stromal and immune cells, creating divergence from *in vivo* signaling environments. As a result, many therapies that show promise in the lab often fail to translate into clinical success.^39^ While 3D-PTAs have shown promise by preserving components of the native TME, reproducibility and standardization are currently major limitations attributable to non-physiologic growth factors/conditions and varying protocols between labs.^4,5^ To address these barriers, we have developed the SMART system: a simple, standardized, reproducible perfusion platform that maintains intact human tumor tissue in its native microenvironment. By using autologous plasma as the perfusate, the SMART system standardizes culture conditions to better approximate *in vivo* biology. With broad implementation, data derived from the SMART system or other physiologically relevant, autologous perfusate cultures will unveil a deep understanding of how the TME responds to a given therapy by allowing for the application of an evolving armamentarium of computational analytics across large numbers of diverse patient populations.

Technological advances in recent years have significantly enhanced our ability to extract robust, high-dimensional datasets from small tissue samples.^1^ This has enabled the integration of multiple layers of molecular data, including genomics, transcriptomics, epigenomics, proteomics, and metabolomics, providing a more comprehensive and systems-level understanding of the TME. While generating large volumes of molecular and phenotypic data is not new, the growing availability of advanced computational tools now allows researchers to uncover patterns and insights that would be difficult to identify through conventional analysis. Importantly, these multi-omic datasets can also be integrated with clinical data, facilitating the discovery of biologically and clinically meaningful correlations. As machine learning and artificial intelligence approaches continue to evolve, the capacity to analyze and synthesize complex datasets will further accelerate cancer research.^40^ In this context, the SMART system represents an ideal preclinical model to fully harness the power of multi-omic analysis and link molecular changes to therapeutic response and clinical outcomes. This manuscript demonstrates the platform’s utility through proof-of-principle experiments focused on drug interrogation. For example, 48-hour exposure to cetuximab in *KRAS* mutant colorectal cancer within the SMART system resulted in increased MAPK, EGFR, PI3K pathway signaling and a shift toward M2 macrophage-associated signaling. While these initial studies highlight individual aspects of the system’s capabilities, its broader potential lies in the ability to systematically interrogate therapeutic responses through the integration of functional data with multi-omic and clinical datasets. Through this method, the SMART system has the potential to provide deeper understanding of how molecularly diverse TMEs respond to similar therapeutics.

The ability to characterize dynamic changes in the TME through multi-omic analysis makes the SMART system particularly well-suited for integration with clinical trial research. To demonstrate this potential, we used bintrafusp alpha, a bifunctional fusion protein combining a PD-L1 antibody with the extracellular domain of TGF-β RIIa, to treat tumor tissue from two patients with HPV-positive metastatic genitourinary cancers. Clinically, one patient experienced short-interval disease stabilization followed by rapid progression, while the other was a clinical responder. Using the SMART system, we interrogated these tumor samples to investigate the discrepancy in clinical response and found clear differences in both immune cell composition and cytokine expression between the two samples. This highlights potential mechanistic underpinnings of response and resistance related to mechanism of action, highlighting the need for further work to understand how best to apply TGF-β inhibition. Notably, the SMART system is compatible with tissue from a core needle biopsy, enabling real-time functional analysis of tumor samples in parallel with clinical trials. Although not designed as a prediction tool, the SMART system provides insight into short-term drug-induced alterations in the TME, enabling the identification of resistance mechanisms and the design of combination therapies tailored to exploit these TME changes to enhance treatment efficacy. Furthermore, obtaining additional tumor biopsies after treatment would allow for comparative analysis of drug-induced changes in the TME between *in vivo* and *ex vivo* contexts, further refining the model and its translational utility.

A significant strength of the SMART system lies in its simplicity, reproducibility, and standardization. It enables the maintenance of *in vivo* parameters through point-of-care instrument evaluations, providing a controlled, physiologic time window to probe tumor biology using a multitude- of cell- and molecular-based technologies. The system accounts for key factors such as shear stress on tissue viability, electrolyte concentrations in the perfusate, physiologic temperature, maintaining pH through the aid of a gas mixer, and carbohydrate metabolism. We demonstrate sustained cell viability across the continuum of cell populations within the TME, including tumor, stromal, and immune cells. Furthermore, this is shown across multiple cancer histologies and tissue/biopsy types, including core needle biopsies and peritoneal metastases. Finally, the use of autologous plasma as the perfusate eliminates the potential cellular alterations related to non-physiologic growth factors/condition inherent to augmented media preparations. This perfusate is easily reproduced in any lab using the system carefully according to detailed standard operation protocols. This validates the platform’s ability to preserve cellular complexity necessary for meaningful interrogation and its user-friendly, operationally straightforward design.

The system outlined herein has several limitations. First, it requires tissue from the operating room or interventional radiology suite with minimal delay, obligating coordination between clinical and research teams. During system optimization, we experimented with short-term cold UW storage to extend time between excision and placement in the SMART system, but this substantially altered cell populations. Second, we focused on aggressive intra-abdominal metastases with substantial amounts of tumor-associated fibrosis, which are likely ideal tumors for *ex vivo* study. The results of the system on less aggressive tumors are unknown and must be determined before experimental planning. Next, the system lacks real-time homeostatic feedback that was demonstrated to be essential for long-term (1 week) *ex vivo* perfusion of human livers.^41^ We used point-of-care instruments to make adjustments that require attention at regular intervals. Finally, the use of any *ex vivo* system, including the SMART system, will require the establishment of acceptable “deviation from baseline metrics” in order for experimental data to be collated for detailed computational analyses. Although outside the scope of the current work, we feel that these metrics must be established for each tumor histology, taking into account baseline heterogeneity from the time of sample collection, which can vary greatly in heavily treated patients.

The SMART system represents an advancement for pre-clinical cancer modeling by preserving the native TME through a simple, standardized, reproducible perfusion system. By using autologous plasma and maintaining physiologic conditions, the platform enables functional and multi-omic analyses of intact human tumor tissue while capturing critical interactions between tumor, immune, and stromal cells. Initial proof-of-concept experiments demonstrate its potential to uncover mechanisms of drug action, therapeutic resistance, and inform novel combination strategies. Furthermore, its compatibility with small biopsy samples and integration with clinical trial workflows positions the SMART system as a powerful tool for translational research. Ultimately, this novel pre-clinical model will accelerate efforts that aim to improve efficacy of cancer treatments.

## Acknowledgments

This research was supported by the Intramural Research Program of the National Institutes of Health (NIH) and with Federal funds from the National Cancer Institute, National Institute of Health, under Contract No. HHSN26120150003I. The contributions of the NIH author(s) are considered Works of the United States Government. The findings and conclusions presented in this paper are those of the authors and do not necessarily reflect the views of the NIH or the U.S. Department of Health and Human Services. JMH had full access to all the data in the study and takes responsibility for the integrity of the data and the accuracy of the data analysis. We acknowledge with gratitude the efforts of the patients and their families for their participation and support of these clinical trials. Bintrafusp alfa was provided by EMD Serono (CrossRef Funder ID: 10.13039/100004755). Flow cytometry acquisition were performed at the CCR LGI Flow Cytometry Core supported by funds from the Center for Cancer Research.

## Star Methods

### Resource availability

#### Lead contact

Further information and requests should be directed to and will be fulfilled by the Lead Contact, Jonathan Hernandez (Head of Liver Metastasis Biology Section, Surgical Oncology Program, Center for Cancer Research, National Cancer Institute, National Institutes of Health).

#### Materials availability

This study generated one unique reagent, a pan-anti-MHC-I chimeric antibody, W6/32LALAPG, which binds to the overlapping a3b2 site of MHC-I.

#### Data and Code availability

Raw data will be deposited at EGA and will be made available upon reasonable request upon submission to the final publisher.

### Experimental model and subject details

Patients undergoing abdominal surgery for germline mutations or metastatic tumors were consented for tissue and blood donation under IRB-approved protocol NIH 13-C-0176l. For the purposes of this project, intraoperative biospecimens were collected from 129 patients over a six-year period totaling over 150 smart systems with a mean duration of 82.6 hours (**Supp Table 1**). Overall, normal peritoneum system optimization was conducted in 49 (38.0%) of the system runs. Peritoneal tumor deposits composed 53 (41.1%) of the system runs while tumor sectioning either via thin sectioning or core biopsies accounting for 27 (20.9%). Heparinized whole blood was collected from each patient under sterile conditions and released for research purposes after fulfilling all clinical requirements. Sterile specimens from abdominal surgeries were processed immediately upon resection under the direction of the surgeons (JMH, JLD, AMB) and in consultation with Pathology staff (DK). To maintain maximal sterility of the specimens, the latter were placed in a sterile field within the operating room. Following the directives of the pathology representative tissue that was not needed for diagnostic evaluation was procured in a sterile environment immediately upon resection.

### Method Details

#### Tissue Procurement

Immediately after surgical resection, excess peritoneal tissue not needed for pathological diagnosis and < 0.5 mm in thickness was aseptically placed in warm DMEM (Gibco, Grand Island, NY, USA) supplemented with anti-anti (Gibco, Grand Island, NY, USA). Following visual inspection of the resected peritoneal tumor deposits and normal peritoneum, excess adipose tissue is stripped carefully, and the peritoneum is draped over the small ring of SMART platform with the mesothelial surface facing outward. For tumor bearing peritoneum, macroscopically visible tumor is positioned over the center of the platform before securing it in place. Samples were affixed onto autoclaved 3D printed custom platforms using 2-0 silk suture and were trimmed of excess tissue. Platforms were inverted into 24-well culture dishes containing perfusate, transferred immediately to the laboratory, placed in the pre-oxygenated system. Control tissues were fixed in 10% formalin, snap frozen, or dissociated based on experimental plans. Following the SMART system run the same process was also performed. Perfusate samples were snap frozen for further analyses as necessary. The duration of time to affix the tissue to a platform is an average of 45 seconds. Depending on the experimental set up, tissue was affixed to either regular platforms or platforms with large openings (**Figure 1D**).

Tumors that were not peritoneal metastases were processed by sectioning or core sampling. Sections for mounting onto the SMART platforms were prepared using a Compresstome vibratome. In brief, following procurement tissue biopsies were cut into pieces of approximately 1.5 x 1.5 cm, secured to the specimen tube base with glue and embedded in prewarmed 2.5% agarose. Agarose solidification was facilitated using a chilling block and serial sections were cut using the following parameters: thickness 300 um, advance speed 4 and oscillation rate 5-6. Sections were collected in warm DMEM (Gibco, Grand Island, NY, USA) supplemented with anti-anti (Gibco, Grand Island, NY, USA). Core biopsies were taken directly from procured tissue biopsies using a 18 gauge core biopsy needle. Both, slices and cores were immediately mounted on a platform using a 2-0 silk suture and transferred to a pre-oxygenated system. For slices, regular platforms with round opening were used whereas core biopsies were affixed to grooved biopsy platforms (**Figure 1D)**, that can receive up to 4 tissue pieces.

#### Perfusate Preparation

The perfusate is composed primarily of autologous patient plasma, which is drawn as whole blood from the patient preoperatively and spun down to allow for separation and removal of the red cells and buffy coat. The perfusate is composed of autologous plasma. Insulin (0.1 mL per 5), 5% dextrose (0.1 mL per 5 mL), and Anti-Anti (Pen-Strep plus Amphotericin, Gibco, Grand Island, NY, USA, 0.05 mL per 5 mL) are added to the plasma to create the final perfusate. The perfusate is prepared in a sterile fashion inside a biosafety cabinet. Each SMART System requires 13 milliliters of perfusate to fill the chamber and circuit.

#### SMART system setup

The SMART perfusion circuit (PCT Patent Application No. PCT/US2021/021525) allows the maintenance of *ex vivo* viability of tissue for drug testing, mechanistic interrogations, and to study temporal effects at near-physiologic conditions.

The circuit is assembled from SMART custom 3D printed components, an oxygenator connected to a gas blender, a peristaltic pump, a sampling port, and a free water infusion line, all connected with sterile tubing as depicted in **Figure 1B**. 3D printed components were autoclaved before and between uses. To maintain sterility, 3D printed parts, oxygenator and sample port were connected with tubing in a biosafety cabinet. The peristaltic pump was housed in a 37 °C incubator and can be used to supply 2 circuits at a time.

Before introducing tissue, the system is primed/optimized. To this end, perfusate is added to the circuit and equilibrated to 37 °C. A humidified gas flow through the oxygenator is established at 80 milliliters/smart system/minute. The gas mix is composed of oxygen and carbon dioxide, blended in variable concentrations via a gas mixer; optimal starting gas mix was defined in **Figure 1E**. The pH of the perfusate was regulated by adjusting the amount of carbon dioxide in the gas mix.

To maintain physiological parameters following tissue placement into the platform holder, point of care testing was carried out at set time intervals of approximately every 4-6 hours with an analyzer dependent on the behavior of the system (Nova Prime+, Nova Biomedical, Waltham, MA, USA). pH, oxygen levels, and electrolyte concentration were assessed. To compensate for evaporative loss of water, continuous infusion of sterile water was initiated using a metered syringe injector. Indicated adjustments were made by changing the gas mix, increasing or decreasing the free water infusion rate, and adding supplements such as dextrose or sodium bicarbonate. Every 48 hours, a complete perfusate exchange was performed to remove cellular waste and refresh innate plasma components for runs that exceeded 48 hours. A list of SMART system components and reagents are provided in **Supp Table 2**.

Experimental drugs were introduced into the perfusate as indicated by the experimental design. Tissue samples taken immediately following procurement were preserved in 10% neutral buffered formalin and served as a control (Time 0) for a given SMART experiment. Throughout the experiment, SMART platforms and/or perfusate samples were removed at indicated time points and subjected to analysis. At the end of each SMART run one tissue sample was preserved in 10% neutral buffered formalin, embedded in paraffin and cut at 5 μm. Hematoxylin and Eosin (H&E) stained slides were evaluated by a pathologist. Perfusate samples collected throughout an experiment were snap frozen at time of collection and analyzed together at the end of an experiment to avoid batch effects. Further tissue analysis is detailed in the method sections below.

#### Histology and Immunohistochemistry Analysis

A list of antibodies used in this study including vendor and clone information is provided in **Supp Table 3**.

##### H&E, lowplexIF and IHC

###### Hematoxylin and Eosin (H&E) Staining

Tissues were formalin-fixed, embedded in paraffin and sectioned at 5 μm thickness. Routine hematoxylin and eosin (H&E) staining was performed using the Sakura Tissue-Tek Prisma™ automated stainer. The slides were dewaxed using xylene and then hydrated using a series of graded ethyl alcohols and stained with commercially available hematoxylin, clarifier, bluing reagent, and eosin Y. A regressive staining protocol was employed, in which tissues are initially overstained and then subjected to differentiation using clarifier and bluing reagents to selectively remove excess stain. Following staining, slides were cover-slipped using the Sakura Tissue-Tek Glas™ automated cover slipper and allowed to dry prior to microscopic evaluation.

###### Masson’s Trichrome Staining

FFPE slides were deparaffinized and hydrated to distilled water. Slides were then mordanted in Bouin’s solution for 1 hour at 60°C. Following mordanting, sections were stained with Biebrich Scarlet, which binds to acidophilic tissue elements such as muscle, collagen, and cytoplasm. Subsequently, slides were treated with phosphotungstic and phosphomolybdic acids to selectively remove Biebrich Scarlet from collagen, while retaining it in cytoplasmic elements. Collagen was then visualized by staining with Aniline Blue. After staining, the sections were dehydrated through graded alcohols, cleared in xylene, and coverslipped with Paramount mounting medium. Slides were then ready for imaging and histological review.

###### Single Stains-Automated

Single IHC stains were performed on the Bond RX autostainer (Leica Biosystems) using the Bond Polymer Refine Kit (Leica Biosystems DS9800) and the following conditions: Epitope Retrieval 1 (Citrate) for CD68, Cleaved Caspase 3, and CD3, Epitope Retrieval 2 (EDTA) for CD4, PD-L1, and CD44, secondary antibody Rabbit anti-Rat IgG for CD3. Isotype control reagent was used in place of primary antibodies for the negative controls. Slides were removed from the Bond autostainer, dehydrated through ethanols, cleared with xylenes, and coverslipped.

###### Single Stains-Manual

Single IHC staining for E-Cadherin was performed manually using Citrate antigen retrieval, Biotinylated Goat anti-Rabbit IgG (Vector Labs), ABC-HRP (Vector Labs), and DAB. Sections were counterstained with hematoxylin, dehydrated through ethanols, cleared with xylenes, and coverslipped.

###### Double Fluorescent Stains-Automated

Double fluorescent sequential stains performed on the Bond RX autostainer (Leica Biosystems) used the Bond Polymer Refine Kit (Leica Biosystems DS9800), with omission of the PostPrimary reagent, DAB and Hematoxylin. For CD45/Ki67, after antigen retrieval with Citrate (Bond Epitope Retrieval 1) sections were incubated for 30 min with CD45, followed by the Bond Polymer reagent and OPAL Fluorophore 520 (AKOYA). The CD45 antibody complex was stripped by heating with Citrate. Sections were then incubated 30 min with Ki67, followed by the Bond Polymer reagent and OPAL Fluorophore 690. For NCAM1/CD69, after antigen retrieval with EDTA (Bond Epitope Retrieval 2) sections were incubated for 30’ with NCAM1, followed by the Bond Polymer reagent and OPAL Fluorophore 520 (AKOYA). The NCAM1 antibody complex was stripped by heating with EDTA. Sections were then incubated 30 min with CD69, followed by the Bond Polymer reagent and OPAL Fluorophore 690. For CD69/CD8, after antigen retrieval with EDTA (Bond Epitope Retrieval 2) sections were incubated for 30 min with CD69, followed by the Bond Polymer reagent and OPAL Fluorophore 690 (AKOYA). The CD69 antibody complex was stripped by heating with EDTA. Sections were then incubated 30 min with CD8, followed by the Bond Polymer reagent and OPAL Fluorophore 520. Slides were removed from the autostainer, stained with DAPI and coverslipped with Prolong Gold AntiFade Reagent (Invitrogen #P36930). Images were captured using the AKOYA PhenoImager whole slide scanner.

###### Double Stains-Manual

Double fluorescent stain was performed manually. After antigen retrieval with EDTA (Agilent #S2367) sections were incubated overnight at 4°C with pSTAT3, followed by HRP conjugated anti-rabbit IgG and OPAL Fluorophore 520 (AKOYA). The pSTAT3 antibody complex was stripped by heating with Citrate (Vector Labs). Sections were then incubated overnight at 4C with pS6, followed by HRP conjugated anti-rabbit IgG and OPAL Fluorophore 690 (AKOYA). Double fluorescent stain (phospho-AKT/phospho p44/42 MAPK) was performed manually. After antigen retrieval with Citrate (Vector Labs) sections were incubated overnight at 4°C with pAKT, followed by HRP conjugated anti-rabbit IgG and OPAL Fluorophore 520 (AKOYA). The pAKT antibody complex was stripped by heating with Citrate. Sections were then incubated overnight at 4°C with p44/42 MAPK, followed by HRP conjugated anti-rabbit IgG and OPAL Fluorophore 690 (AKOYA). Sections were stained with DAPI and coverslipped with Prolong Gold AntiFade Reagent (Invitrogen). Images were captured using the AKOYA PhenoImager whole slide scanner.

#### Multiplex IF staining using Ultivue 12-plex assay

The in-situplex multiplex immunofluorescent assay, commercialized by Ultivue, enables detection of multiple protein targets in tissues by target amplification and cyclic staining/signal removal. FFPE tissue sections were stained using Ultivue’s 12-plex panel with antibodies for selected immunomarkers and PanCK/SOX2 for detection of cancer cells. The 12-plex panel consisted of antibodies for CD3, CD4, CD8, CD20, CD56, CD68, CD163 to detect T cells, B cells, NK cells, M1 and M2 macrophages, FoxP3, Granzyme B, PD-1 and PD-L1 for their functional state and expression of immune checkpoint markers. Tissues were baked overnight at 60°C to improve tissue attachment. Dewax and antigen retrieval was performed on Leica Bond Rx using ER2 antigen retrieval with the recommended protocol from the vendor. Slides were stained manually following Ultivue’s recommended protocol for blocking, antibody staining, signal amplification, nuclear staining and signal visualization followed by mounting and coverslipping. The 12 signals were processed in three cycles and imaged on an Axioscan 7 slide scanner set to detect signals on FITC, TRITC, CY5 and CY7 channels + DAPI. After imaging, the coverslips were removed by soaking in PBS overnight followed by signal removal and detection of targets in the next cycle. After the last cycle, the signals were removed, and the tissue was stained for H&E using Parhelia Spatial Station (Parhelia Bio) followed by imaging of Axioscan 7.

The whole slide images from three rounds and the H&E staining were registered and fused into a single composite image using UltiStacker.AI cloud-based software platform, provided by Ultivue as a service. The composite images were loaded into HALO (NCI-HALO version 3.6, a centrally supported cloud-based enterprise solution for quantitative image analysis) for viewing and analysis.

#### CODEX: Phenocycler fusion assay for multiplex protein detection

For the Phenocycler Fusion multiplex protein detection assay, slides containing FFPE sections (4 um) were processed and stained with an antibody cocktail containing a mix of commercially available and custom conjugated antibodies following recommendations from Akoya Biosciences and previously described methods.^42^

Slides were baked overnight at 60°C to improve attachment of tissues to glass. They were immersed in xylene (2 times, 10 min) to remove paraffin and later rinsed in a decreasing alcohol series to rehydrate the tissues (100%, 100%, 90%, 70%, 50%, 30% ethanol in MilliQ water for 4 min each). Antigen retrieval was performed in a pressure cooker set to low pressure using antigen retrieval solution pH 9 for 15 min (AR9 from Akoya Biosciences). After the slides cooled down to RT, they were soaked in Hydration (2 times, 2 min) and Staining buffer (up to 20 min). Antibodies were combined as indicated in staining buffer containing J, N, S and G blocking reagents (Akoya Biosciences) and were added to tissues overnight at 4°C. The slides were washed in staining buffer (2 times, 2 min) and the bound antibodies were fixed using 1.6% paraformaldehyde (diluted in storage buffer) for 10 min. The slides were washed in PBS (3 times, 2 min), incubated in ice cold methanol for 5 min and, after additional PBS washes (3 times, 2 min), were incubated in the crosslinker solution diluted in PBS for 20 min. After additional PBS washes (3 times, 2 min), the stained slides were imaged on Phenocycler fusion or stored at 4°C in storage buffer for no more than 2 weeks. For imaging, a flow cell was created on the slide and the imaging run was set-up following the vendor’s recommendations. Details about antibody clones and dilutions are listed in **Supp Table 3**.

At the end of the run, the generated 8-bit images were further processed to generate 16-bit images to reduce signal saturation on selected channels. Both the 8 and 16-bit images were uploaded into HALO image analysis software (NCI-HALO version 3.6) for visualization and analysis.

##### Post-CODEX H&E staining

After the CODEX run finished, the flow cell was disassembled by soaking it in Xylene for at least 3 hours. The slides were rinsed into the following diluent series: ethanol diluted in MilliQ water (100%, 100%, 70%, 30%) for 3 min each and PBS (3 times, 3 min). H&E staining was performed using the Parhelia Spatial Station autostainer (Parhelia Bio). The sections were mounted using Permount mounting medium (Fisher Scientific SP15-500) and scanned using a Zeiss Axioscanner 7.

##### Antibody-conjugation with oligonucleotide-tags

Carrier free antibodies were conjugated to selected barcodes using commercial reagents following Akoya Bioscience’s recommended protocols as described earlier.^43^ First, preservatives and other additives (like trehalose) were removed by using Amicon Ultra 30K centrifugation filters (Millipore, Cat. No. UFC503024). After multiple washes with PBS, the protein concentration was measured using an Implen nanophotometer with IgG mouse settings. 50 ug of protein was used for conjugation, diluted in 100 ul PBS. For conjugation, Amicon Ultra 50K centrifugation filters (Millipore, Cat. No. UFC505024) (one for each conjugation) were washed with 500 ul filter blocking solution (Akoya Biosciences, Cat. No. 7000009 Part No 232113) and centrifuged at 12,000 x g for 2 min. The remaining solution was discarded and 50 ug antibody in a volume of 100 ul supplemented with PBS was added to the filter followed by centrifugation at 12,000 x g for 8 min. 260 ul antibody reduction master mix (Akoya Biosciences, Cat. No. 7000009 Part No. 232114 and Part No.232115) was added to the filter and incubated for 30 min. After centrifugation at 12,000 x g for 8 min, the filter was washed with 450 ul of conjugation buffer (Akoya Biosciences, Cat. No. 7000009 Part No. 232116). The barcode (one vial for each antibody) was resuspended in 10 ul nuclease free water (Ambion AM9938) and complemented with 210 ul of conjugation buffer. The mix was added to the filter and incubated for 2 hours. The filter was centrifuged at 12,000 x g for 8 min followed by washes (three times) with 450 ul purification solution (Akoya Biosciences, Cat. No. 7000009 Part No. 232117) and resuspended into 100 ul of antibody storage solution (Akoya Biosciences, Cat. No. 7000009 Part No. 232118). The antibody was collected by inverting the filter and centrifuging the content into a new tube at 3000 x g for 2 min. The collected conjugated antibody was stored at 4°C.

#### MxIF Analysis

##### QC Step

MxIF data included either 25 or 52 antibodies. Each marker expression was assessed by a pathology analyst to verify signal quality, specificity, localization (membrane, cytoplasmic, or nuclear), intensity and background noise. Markers were classified as “Failed”, “Borderline”, “Passed” based on the following criteria:

● Passed: Specific signal (nuclear, membrane, or cytoplasmic) with low background.
● Borderline: Specific with visible background: Artifacts might be present, but not affecting the analysis.
● Failed: Non-specific staining, extensive artifacts, high background, or absent signal.

For each “Passed” and “Borderline” markers, thresholds were set to differentiate true cell staining and background, reducing false positive expression read and improving clustering quality.

##### Artifact Detection

Artifact detection was integrated into preprocessing to identify and exclude regions with technical artifacts. A semantic segmentation model utilizing UNet++ architecture with EfficientNet-B6 encoder was implemented for precise artifact localization. The model was trained on a dataset containing common artifacts, including blurring, tissue folds, biomarker edge staining, and overstained regions. By systematically excluding these regions, the minimized erroneous signal interpretation, improving the accuracy of downstream analysis.

##### Cell Segmentation

Cell segmentation was performed using a UNet++ semantic segmentation neural network with MaxViT backbone. In each patient’s IF image, DRAQ was designated as the primary nuclei marker due to its specificity for nuclear material. The primary membrane marker was determined by combining multiple membrane/cytoplasmic markers, optimized per patient based on signal quality to ensure accurate membrane delineation. The segmentation model was pretrained on a manually annotated internal dataset, consisting of images with three distinct channels: nuclear, membrane, and a composite membrane/cytoplasmic channel. The composite channel was generated by selecting the brightest pixel across all membrane/cytoplasmic channels at each pixel position. Each segmented cell was assigned a unique identifier with spatial coordinates. Marker expression was quantified per cell by measuring mean signal intensity and marker-positive area coverage.

##### Cell Typing

Following quality control by a pathology analyst, a set of usable markers was selected, intensity signal thresholds were applied to exclude artificial background staining. Each cell was assigned a unique identification number and key metrics were calculated including signal (mean intensity and marker+ area per cell) and spatial metrics (position of centroid on x and y axis). By utilizing the area of each marker expression for every cell, each cell was characterized based on marker expression within its contour, by utilizing the area of each marker expression. Leiden clustering was then applied to classify major cell populations marker expression areas. The quality and consistency of these populations were assessed by a pathology analyst before further in-depth analysis.

##### Cell Typing of Subpopulations

Following the identification of major cell populations, cell subtypes within immune, stromal and tumor populations were assessed marker expression patterns. Key markers included to identify immune cells were Ki-67, PD-1, PD-L1, Granzyme B, and HLA-while HER2, Ki67, PCNA, PR, and Fibronectin were used for tumor cells. Cell subtyping was performed using mean intensity and marker^+^ area to enhance classification accuracy. For low intensity markers, a gating strategy was applied by setting an intensity threshold to determine marker positivity. Validation of subtype assignment was performed through spatial mapping, where cells were overlaid onto the original MxIF image and color-coded according to subtype. Additionally, for markers PD-1, PD-L1, HLA-DR, Ki-67, PCNA, Granzyme B, EPCAM, E-cadherin, B-catenin, IFNG, CD44, BCL-2, TP63, and HLA-A the percentage of positive cells within each population were quantified to assess expression patterns across control and treatment groups over time.

##### Composition of cell populations in Low-Plex images

Due to the limited marker set in Low-Plex images and variability across samples, the fraction of tissue with positive expression was calculated. The absence or low-quality of membrane markers impaired segmentation accuracy, while dense nuclear staining often obscured nuclear contours leading to incorrect contour approximations. Marker coverage was assessed by intersecting the nuclear marker (DAPI) with each present marker, enabling the calculation of the fraction of positive cells area while excluding tissue regions devoid of cells.

##### Additional quantification and metrics

For patients with tumor and tissue masks, cell populations within and outside the tumor were quantified as percentage. For CODEX patients, cell density was calculated as the number of cells per mm^2^ of tissue. For Patient 11, subsamples were selected to account for intra-group heterogeneity.

##### Dextran uptake assay

Tissue platforms at 0 and 96 hr were incubated in 1ml of prewarmed perfusate containing 5ul CF595 dextran (5 mg/ml in PBS, Biotium 80114), 7.5 ul of AlexaFluor488 anti-human HLADR antibody (Biolegend 307620, Clone L243) and 2 drops of Nuc Blue (Thermo Fisher R37605) in the incubator for 2 hr. Platforms were removed from the perfusate, briefly washed with PBS and fixed for 24 hr in 1% paraformaldehyde in PBS. Following 2 PBS washes, samples were mounted on an imaging adaptor^44^ and imaged at a Leica Stellaris white light laser confocal microscope using a 20x objective. 1-3 12 um 9-tile Z-stack scans per platform were acquired. Maximum projections were analyzed in HALO using the Cytonuclear FL analysis module (v2.0.12). The percentage of cells positive for both, CF595 dextran and AlexaFluor488 HLA-DR antibody of all cells positive for AlexaFluor488 HLA-DR only were used to determine the percentage of cells actively taking up dextran.

##### Cloning and Generation of purified W6/32LALAPG Antibody

The W6/32 monoclonal antibody was purchased from BioXcell. DNA sequences of the VH, VL and Fc segment of W6/32 were identified by sequencing. The mouse Fc segment of W6/32 was subcloned into pFUSE-CHIg-hG1 (Invivogen, San Diego, CA) to replace with human IgG1 Fc to generate W6/32-hIgG1. To prevent the binding of Fc segment of W6/32-hIgG1 to FC gamma receptors (FcgR) on human immune cells, L234A, L235A, and P329G mutations were introduced in the H chain constant region by site-directed mutagenesis with QuikChange Lightning Site-Directed Kit (Agilent, Santa Clara, CA, USA) following manufacturer’s instructions to generate W6/32LALAPG. Plasmids encoding the W6/32 H and L chains were mixed at a ratio of 1:2 respectively and transfected in 293F cells with lipofectamine (ThermoFisher Gibco, USA). After 6 days the culture supernatant containing the secreted W6/32LALAPG antibody was harvested and antibody purified by affinity chromatography using Protein A Sepharose (Cytiva, Uppsala, Sweden).

##### Bulk RNA Sequencing

RNA was extracted from snap frozen tissue and formalin embedded tissue. Bulk RNA sequencing TruSeq Stranded Total RNA (Illumina) was used for RNA library construction. The libraries were sequenced on a NovaSeq S4 with 100M reads/sample and a sequencing depth of 100 M reads (2 × 150 pairs). Samples were pooled and sequenced on NovaSeq 600 using Illumina(R) Stranded Total RNA prep, Ligation with Ribo-Zero Plus and paired-end sequencing.

##### RNA-seq data processing, quality control, and analysis

Quality control of fastq files was performed using FastQC (v0.11.9),^45^ FastQ Screen (v0.14.0),^46^ RSeQC (v3.0.1),^47^ and compiled with MultiQC (v1.13).^48^

PathSeq was used to identify microbial organisms. HLA comparison, and SNP coverage distribution in matched samples were used to validate sample correspondence. RNA-seq reads were aligned using Kallisto (v0.43.0)^49^ to the Ensembl GRCh37 reference transcriptome. Noncoding RNA, mitochondrial transcripts, histone-related transcripts and short TCR/BCR transcripts were removed, resulting in 20,062 transcripts. Gene expressions were normalized as transcripts per million (TPM) and log2-transformed.

##### Cell deconvolution

From bulk RNA-seq data was performed using Kassandra algorithm,^16^ predicting the fractions of CD4+ and CD8+ T cells, B cells, macrophages, neutrophils, fibroblasts, endothelial cells and additional subpopulations. Log2-fold changes in cell proportions were calculated comparing Day 2/4 vs. Day 0 samples, and Treated Day 2 vs. Untreated Day 2 samples.

##### Differential Expression Analysis

Reads used FeatureCounts (v. 1.0.1)^50^ for and PyDESeq2 (v0.4.0).^51^ Differentially expressed genes (DEGs) were defined as adjusted p-value <0.005” and log2 fold-change >2. DEGs were used for Functional enrichment analysis of DEGs was assessed using the cancer hallmark gene set from the MSigDB database^52^ using Python-based version decoupler package (v. 1.6.0), integrated with OmniPath.^53^

##### Gene Signature Score Calculations

Single sample gene set enrichment analysis (ssGSEA) scores were calculated using an in-house Python implementation^54^ and MAD-scaled within the cohort.

##### Single cell Sequencing

Tissue was dissociated by mincing with scissors in dissociation media (1X HBSS (Corning) containing 2% bovine serum albumin (Sigma), 50 µg/mL Liberase TM (Roche), and 500 µg/mL DNase I (Sigma Aldrich)), and processing in a GentleMACS OctoDissociator (Miltenyi Biotec) according to the manufacturer’s program h_tumor_02 2 times, followed by incubation in a 37°C shaking water bath at 100 rpm for 30 min. Disrupted tissue was subsequently passed through 70 µm cell strainers (Falcon), followed by ACK lysis, washed in MACS Wash supplemented with bovine serum albumin (Miltenyi), and passed through 40 µm cell strainers (Falcon). Once single cell suspensions were obtained the cell concentration and viability were assessed using the Luna-STEM fluorescent cell counter. Samples that passed initial quality checks were loaded in the lanes according to the 10X Chromium Single Cell 3’ User Guide, with one capture lane per sample targeting recovery of 6,000 cells per lane. Reverse transcription and barcoding were performed immediately. All subsequent steps of library preparation and quality control were performed as described in the 10X User Guide for Single Cell 3’. Sequencing was performed by SCAF in Building 41 in Bethesda utilizing one NovaSeq run. Base calling was performed using RTA 3.4.4, demultiplexing was performed using cellranger v7.0.1 (Bcl2fastq 2.20.0), and alignment was performed using cellranger v7.0.1 (STAR 2.7.2a). Sequenced reads were aligned to the 10x Genomics provided human reference sequence (refdata-gex-GRCh38-2020-A).

##### scRNA-seq quality control and data correction

The raw scRNA-seq data were processed using CellRanger (v) with the human genome as the reference to generate a UMI count matrix. The aggregated CellRanger output files for both SMART system and organoid experiment were converted into the AnnData object using python 3.10 and scanpy v1.9.5.^55^ Cells with >30%mitochondrial gene content, <300 detected genes or predicted as doublets using Scrublet v0.2.3^56^ were digitally filtered out. UMI counts were normalized to 10,000 counts per cell and log-transformed. Cell cycle phase was assigned for using Scanpy function “score_genes_cell_cycle”, and clusters dominated by G2M phase or S phase were excluded.

##### scRNA-seq dimensionality reduction and clustering

PCA was performed using highly variable genes (n=3000) identified using the *scanpy* function “*highly_variable_genes*”. The data were then scaled, analyzed for principal components, and visualized using UMAP. The local neighborhood size was set to 20 nearest neighbours, which also served as the basis for partitioning the dataset into clusters using the Leiden community detection algorithm. A range of resolutions (0.1–1) was utilized to establish a sufficient number of clusters to separate known populations based on expression of established marker expression.

##### scRNA-seq dataset heterogeneity and cell annotation

Cell clusters were annotated based on marker gene expression: tumor epithelial cells (KRT19, EPCAM), monocytes/macrophages (FCGR3A, CD163, PECAM1), lymphocytes (PTPRC, CD3E), Tregs (PTPRC, FOXP3), Fibroblasts (COL1A1). Tumor epithelial cell heterogeneity was assessed using infercnvpy v0.4.4 [https://github.com/broadinstitute/inferCNV] on full unscaled expression data from SMART system and organoid experiments separately. Lymphocytes and monocytes/macrophages served as reference normal cells for CNV inference. PCA followed by neighborhood detection and Leiden clustering with default parameters were performed using inferred CNV scores. Epithelial tumor cells in the SMART system dataset were classified into two groups based on CNV accumulation scores: tumor clone and tumor subclone. To associate mutations with tumor cell clusters BAM files from CellRanger processing were parsed to label cell barcodes carrying KRAS G12V mutation.

##### Cell-cell interactions

Cell–cell interactions were inferred using CellPhoneDB^23^ with default parameters and using the database version v4. The interaction strength was estimated based on receptor-ligand expression levels. Significance was assessed using a permutation test (1000 iterations) with normalized gene expression was used as input. Interactions with p value < 0.05 were considered significant.

##### Inference of cell trajectories for tumor clusters

To compare tumor cells from organoids with the two clusters of tumor cells found in SMART system dataset, the data were embedded in diffusion map space instead of PCA space to reduce noise before neighborhood calculation and graph construction. Tumor cell trajectories were inferred using PAGA (partition-based graph abstraction) algorithm^57^ to generalize relationships between clusters.

##### GeoMx Digital Spatial Profiling

Formalin-fixed, paraffin-embedded tissue blocks were cut at 5 μm thickness onto APEX treated slides to improve attachment. After baking at 60°C overnight, deparaffinization and antigen retrieval was performed on Leica BOND RX autostainer using HIER2 (EDTA based antigen retrieval solution for 20 min) and proteinase K (15 min at 37°C 1 ug/ml) following Nanostring’s recommendations. The 5 slides were processed on 2 consecutive days.

RNA probes (NanoString GeoMx Whole Transcriptome Atlas Human RNA probes for NGS, item# 121401102) were added for hybridization at 37°C for 20 hr. The slides were then washed 2 times with stringent wash (50% formamide in 2XSSC for 25 min each) followed by 2 times wash with 2XSSC (2 min each). Slides were blocked with W buffer for 30 min and then incubated with anti-PanCK-AF532, anti-CD45-AF594, anti-aSMA-AF647 and Syto 13 diluted in W buffer for 1 hr. The slides were washed 3 times with 1X SSC (5 min each) and scanned on the GeoMx DSP machine for selection of regions of interest (ROIs), using FITC/525 nm filter to detect nuclei (100 msec), Cy3/568 nm channel to detect PanCK (200 msec), Texas Red/615 nm to detect CD45 (200 msec) and Cy5/666 nm to detect aSMA (200 msec). Based on the morphology of the tissues, ROIs for media, cap, core and normal wall were selected using polygons, with no masking, for a total of 157 ROIs. The combined surface area of ROIs was 24’202347.71 mm^2^ containing 150207 nuclei. Oligonucleotide barcodes from each ROI were UV-photocleaved, collected into 96 well plates, followed by library preparation and sequencing as recommended by the NanoString GeoMx protocol. Sequencing was performed on NextSeq2000 using 2 Standard SBS NextSeq 2000 P3 Reagents (50 Cycles) kits (27 x 27 and index 8 x 8). Raw basecall files were demultiplexed into individual fastqs corresponding to ROI with the illumina DRAGEN BCL convert software version 3.10.12. Fastqs were converted to Digital Count Conversion files using NanoString’s GeoMx NGS Pipeline v. 2.3.3.10.

##### DSP readout and analysis

Digital Count Conversion files, including expression counts, probes, assay metadata, and annotation information were uploaded to the DSP analysis suite for QC and further analysis. Raw GeoMx data were processed using the GeomxTools R package (GeomxTools: NanoString GeoMx Tools. R package version 3.12.1, https://bioconductor.org/packages/GeomxTools) in accordance with the official guidelines. Quality control filtering of segments was performed using the following thresholds: a minimum of 1,000 reads per segment (minSegmentReads = 1000), at least 80% of reads trimmed (percentTrimmed = 80), 80% stitched (percentStitched = 80), 75% aligned (percentAligned = 75), and 50% saturation (percentSaturation = 50). Segments were also required to have a minimum of one negative count (minNegativeCount = 1), fewer than 9,000 NTC counts (maxNTCCount = 9000), at least 20 nuclei (minNuclei = 20), and a minimum area of 1,000 µm² (minArea = 1000). Probe-level filtering was applied by requiring a minimum probe ratio of 0.1 (minProbeRatio = 0.1) and no more than 20% of probes failing Grubbs’ test (percentFailGrubbs = 20); negative probes were excluded. After filtering, the resulting dataset comprised 147 samples and 13,891 genes.

Downstream analyses were conducted using the decoupler Python package (Decoupler: A Python package for enrichment statistical methods. GitHub repository. Available at: https://github.com/niklasbinder/decoupler-py). Differential expression analyses were performed separately within each patient, comparing pre-treatment and post-treatment samples. Genes that exhibited opposing expression dynamics between patients were classified as responder (R)-specific or non-responder (NR)-specific. For bulk RNA-seq data, log fold changes were calculated as the difference in mean logTPM values between post-treatment and pre-treatment samples. Additional analyses, including PROGENy pathway activity inference and gene set enrichment analysis (GSEA) scoring, were performed with decoupler following the developers’ recommended guidelines.

#### Xenium Spatial Transcriptomics Analysis

##### Xenium slide preparation

Sections for Xenium were prepared per Xenium In Situ for FFPE-Tissue Preparation Guide (CG000578 Rev C, 10X Genomics). Briefly, 5 µm sections were cut, floated in an RNAse-free water bath, and carefully placed onto the sample area of a Xenium slide (PN1000465) using a paintbrush. The slides were then dried at room temperature for 30 minutes, followed by a 3 hour incubation at 42°C on a Xenium Thermocycler Adapter plate, atop a C1000 Touch Thermocycler (BioRad) with the lid open. They were then stored overnight at room temperature with a desiccant. The next day, the Xenium slides were processed following the Xenium In Situ for FFPE Deparaffinization and Decrosslinking protocol (CG000580 Rev C, 10X Genomics). In brief, the slides were incubated at 60°C for 2 hours n on the Xenium Thermocycler Adapter plate atop the 96-well block of a C1000 Touch Thermocycler (BioRad) with the lid open cooled, and sequentially immersed in xylene, ethanol, and nuclease-free water to deparaffinization and rehydration. Subsequently, the Xenium slides were assembled into Xenium cassettes (PN-1000566, 10X Genomics), enabling precise temperature control during incubation on the Xenium Thermocycler Adapter plate in a PCR machine with a closed lid. Slides were processed using the “Xenium Slides and Sample Prep Reagents” kit (PN-1000460, 10X Genomics), starting with decrosslinking and permeabilization at 80°C for 30 minutes, followed by a PBS-T wash. Subsequently, the Xenium slides were processed according to the “Xenium In Situ Gene Expression” user guide (CG000582 Rev D, 10X Genomics). Slides were incubated at 50°C for 17 hours with the pre-designed gene expression probe set, “Xenium Human Multi-Tissue and Cancer Panel” (PN-1000626, 10X Genomics), targeting 377 human genes. This was followed by post-hybridization washing at 37°C for 30 min, ligation at 37°C for 2h, and amplification step at 30°C for 2h, and additional washing steps. slides were then treated with an autofluorescence quencher and underwent nuclei staining.

##### Xenium Analyzer setup and data acquisition

Processed Xenium slides, assembled in Xenium cassettes, were subsequently imaged using the Xenium Analyzer, following the guidelines in the “Xenium Analyzer User Guide (CG000584 Rev B, 10X Genomics)”. Run details including Xenium slide IDs and pre-designed gene expression probe set information, was entered into the Analyzer. The instrument was equipped with the necessary consumables: Xenium Decoding Reagent Module A (PN-1000624, 10X Genomics), Xenium Decoding Reagent Module B (PN-1000625, 10X Genomics), Xenium Instrument Wash Buffer (PN-3001198, 10X Genomics), Xenium Sample Wash Buffer A (PN-30001199, 10X Genomics), Xenium Sample Wash Buffer B (PN-30001200, 10X Genomics), Xenium Probe Removal Buffer (PN-30001201, 10X Genomics), Objective Wetting Consumable, extraction tip, and a full rack of pipette tips. Two Xenium slides/cassettes were then loaded into the instrument, initiating the Analyzer’s ‘sample scan’ process, producing images of fluorescent nuclei in each section. These images determined the regions within the scan area to be included in the instrument’s comprehensive scan of the gene expression probe set. For both runs, the entire section was included in the full scan. Post-run, the Xenium slides were carefully removed. Fresh PBS-T was added to each slide/cassette, and the slides/cassettes were covered and stored in the dark at 4°C until subsequent H&E staining. Run data for each slide was copied from the Analyzer onto a Solid State Drive for further in-house analysis.

##### Xenium Data analysis and visualization

Raw FASTQ files and histology images were processed using Xenium Analyzer (10x Genomics Inc.). The cell counts for KRAS WTand KRAS MUT samples were 84,817, 215,462, 167,812, and 232,015, respectively. The median of the number of transcripts per cell was 55, 33, 51, 54, respectively. The number of decoded transcripts per 100 µm² was 39.6, 32.4, 48.3, and 44.6, respectively. Data were analyzed using squidpy and scanpy Python packages. Cells with area <=15 μm^2^ and fever than 25 counts, as well as genes expressed in fewer than 100 cells were filtered. PCA was performed using all genes from the panel. The data were scaled, analyzed for principal components, visualized using UMAP and clustered using Leiden 20 nearest neighbors. Resolutions from 0.1 to1 were tested to optimize clustering of known populations based on marker expression.

##### Cytokine Analysis

Utilizing snap frozen perfusate samples for both treated and untreated SMART runs, Luminex 24-plex and RANTES ELISA assays were performed. Perfusate was collected at 0, 24 and 48 hours with at least quadruplicates for each time point. All numeric cytokine concentration values were cleaned with setting values lower than detection thresholds to 0. For entries reported as ranges, the mean of the range was used to approximate the concentration. Missing values were encoded as NaN and excluded from numerical computations. Temporal cytokine changes were assessed by calculating log -transformed fold changes (LFC) across successive experimental timepoints. For each cytokine, fold changes were computed separately under control and treatment (Bintrafusp alfa) conditions at 0 and 48 hours.

#### Flow Cytometry

##### Isolation and culture of human peripheral blood mononuclear cells (hPBMCs) with FC-silenced anti-pan-MHC-I antibody

Total leukocytes were isolated from 10 ml of human peripheral blood obtained from healthy volunteers at the NIH blood bank or from cancer patients at the National Cancer Institute by centrifuging at 2000 rpm for 15 minutes using Ficoll-Hypaque. Red Blood Cells were removed by ACK lysis buffer and the single-cell suspension of leukocyte buffy coat was washed with FACS buffer. PBMCs isolated from healthy donors or cancer patients were suspended in RPMI 1640 complete medium supplemented with heat inactivated 10% FBS, 2 mM L-glutamine, 1 mM sodium pyruvate, 1 mM HEPES, 0.1 mM nonessential amino acids, 50 mM 2-mercaptoethanol, and 100 U/ml penicillin/streptomycin. Trypan-blue exclusion staining was done to count live cells using a hemocytometer. Following this, 10 mg of W6/32LALAPG or negative control hIgG1LALAPG antibodies were incubated with 0.5X10^6^ PBMCs in 500 ml of medium for 48 to 96 hours.

##### Isolation of tumor infiltrating lymphocytes (TILs)

Core needle biopsies affixed on the SMART platform (B-regular **Figure 1C**) were harvested and dissected into small pieces in complete RPMI medium. Tumor pieces were further dissociated on the Gentle MACS Octo Dissociator (Milteny Biotech) using the 37C_H_TDK_1X3 program. Red blood cells were removed using the ACK lysis buffer and samples filtered through a 70 μm-pore size strainer (BD Falcon) to obtain single-cell suspension. The single-cell suspension was washed additional 2 times with FACS-staining buffer.

##### Flow cytometry

hPBMCs isolated from healthy donors or cancer patients and their corresponding TILs (n=2) were blocked with FCX Trustain (Biolegend) for 5 minutes in 500 ml of complete RPMI-1640 medium. After blocking, cells were washed and stained with specific surface antibodies for 30 minutes at 4°C in FACS staining buffer (1XPBS, 10% heat-inactivated fetal bovine serum). Cells were washed and analyzed in BD LSR Fortessa X-20 cell analyzer and BD FACSDiva Software (version 8.0). Data analysis was further done with Flowjo^TM^V10 (Treestar).

##### Flow cytometry evaluation of tumor immune microenvironment

Tumor-bearing peritoneum was suspended in the SMART system for 48 hr (P-large **Figure 1C**) or 96 hr (P-regular **Figure 1C**) as described above. PBMCs were isolated from buffy coats following layering whole blood over Ficoll-Paque (Cytiva) and washing with 1X PBS (Gibco). Red blood cells were lysed using ACK lysing buffer (K-D Medical). Patient autologous PBMCs were evaluated at day 0 and after 48h or 96h incubation in R10-HS media (RPMI 1640 (Gibco), 10% Heat-Inactivated Human Serum AB (Access Cell Culture), 1X L-glutamine (Gibco)) at 37°C with 5% CO_2_, as fluorescence minus one (FMO) and staining controls. At day 0 and following SMART system incubation, peritoneum was dissociated by mincing with scissors in dissociation media (1X HBSS (Corning) containing 2% bovine serum albumin (Sigma), 50 µg/mL Liberase TM (Roche), and 500 µg/mL DNase I (Sigma Aldrich)), and processing in a GentleMACS OctoDissociator (Miltenyi Biotec) according to the manufacturer’s program h_tumor_02 twice, followed by incubation in a 37°C shaking water bath at 100rpm for 30 min. Disrupted tissue was subsequently passed through 70 µm cell strainers (Falcon), followed by ACK lysis, washing in MACS Wash supplemented with bovine serum albumin (Miltenyi), and passed through 40 µm cell strainers (Falcon).

Cells recovered from dissociated peritoneum and PBMCs were counted and distributed to the wells of a 96-well v-bottom tissue culture treated plate (Corning) according to yield for dissociated peritoneum (∼230k-380k cells per well 96h SMART, ∼550k-975k cells per well 48 hr SMART) or 1 million cells per well for PBMC staining controls. For 96 hr SMART, cells were stimulated for 3h in the presence of 10 µg/mL Brefeldin A (Sigma Aldrich) at 37°C with 5% CO_2_ and then kept at 4°C overnight prior to staining. Wells were either unstimulated or were stimulated with 1 µg/mL staphylococcal enterotoxin B (SEB, Calbiochem) or with 50 ng/mL phorbol-12-myristate 13-acetate (PMA) and 1 µg/mL ionomycin (Cell Activation Cocktail without Brefeldin A, Biolegend). FMO controls were stimulated with 25 µg/mL ImmunoCult™ Human CD3/CD28 T Cell Activator (StemCell Technologies). For 48h SMART, dissociated cells were plated and kept at 4°C overnight prior to staining, without prior stimulation.

Cells were sequentially stained with amine reactive viability dye, extracellular antibody cocktail, permeabilized with Foxp3 Transcription Factor Fixation/Permeabilization reagents (eBioscience), and subsequently stained with intracellular antibody cocktail. Both extracellular and intracellular antibody cocktails were prepared using pre-titrated antibody quantities with BD Horizon Brilliant Stain Buffer Plus (BD Biosciences) and were stained and washed with FACS Buffer (1X PBS (Gibco), 1% bovine serum albumin (Sigma), and 0.1% sodium azide (Sigma Aldrich)) consistent with established procedures. Fixed stained cells were kept at 4°C overnight prior to acquisition on a BD Symphony cytometer, and data were analyzed using FlowJo v 10.10.0 (BD Life Sciences) (ref). Gating was validated by FMO controls. Dissociation, stimulation, staining, and acquisition were performed both at baseline and at the experiment endpoint (48 hr or 96 hr) as applicable for each experiment.

---

**Figure S1.**
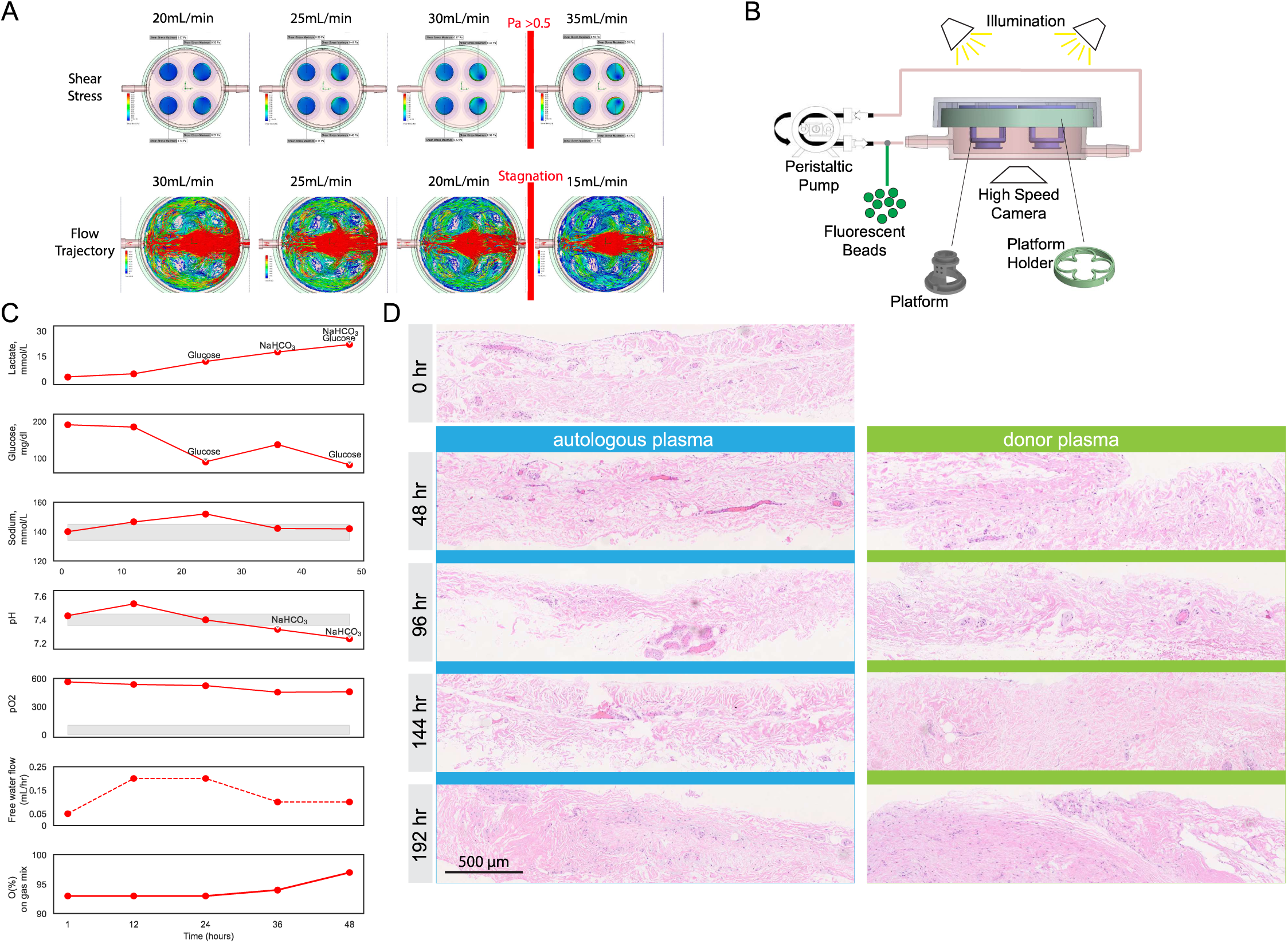
**A:** Additional data points to determine the effect of flow rate on shear stress and flow trajectory within the perfusion circuit as presented in Figure 1D. **B:** Set up for the iterative angle change experiment. **C:** Changes in lactate, sodium, and glucose concentrations and pH ranges for an additional patient. **D:** Set of H&E stained FFPE sections from tissue used to quantify nuclear loss using autologous versus donor plasma for the generation of perfusate (Figure 1 **I,J**).

**Figure S2.**
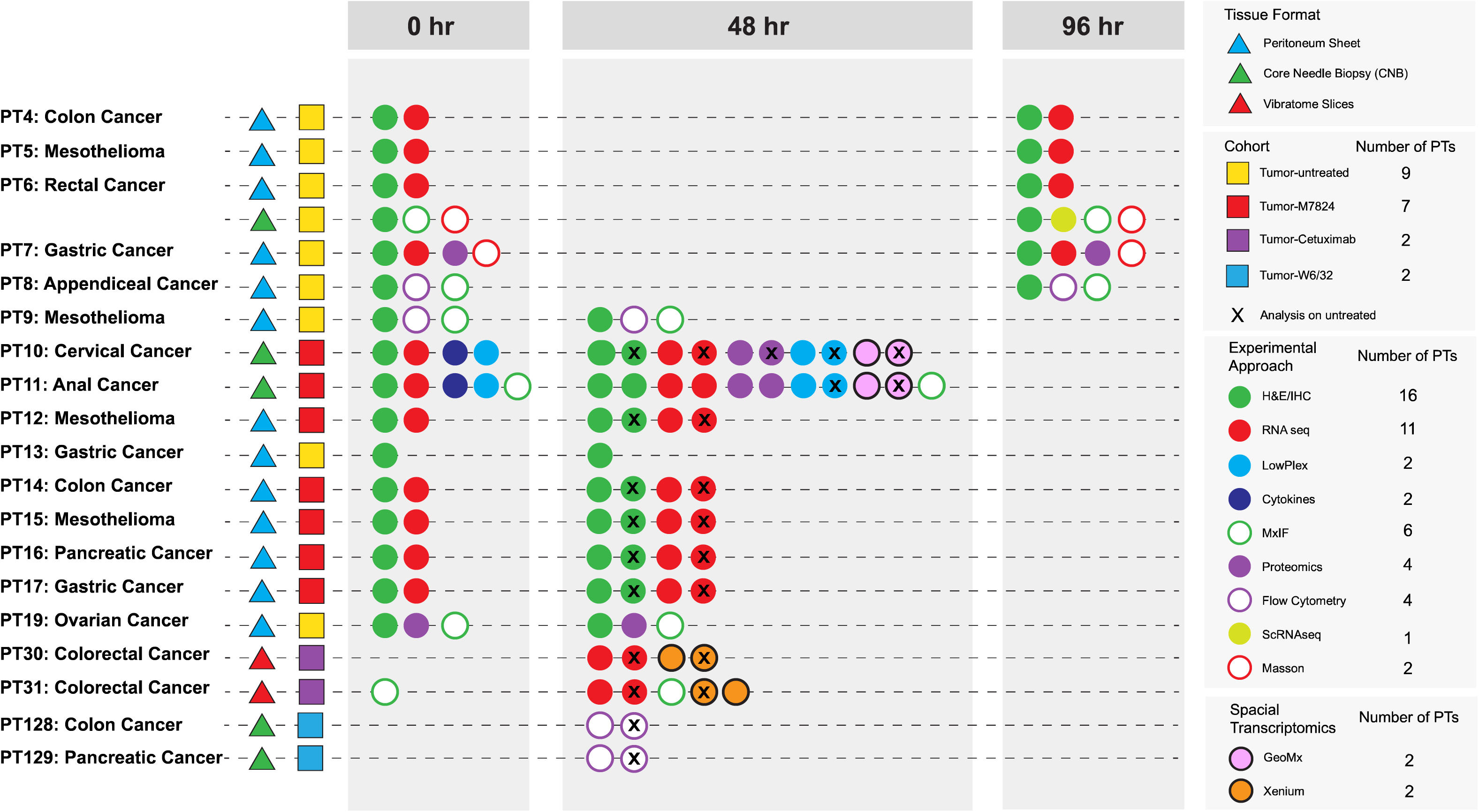
Summary of cohort composition and interrogation techniques applied. PT=Patient

**Figure S3.**
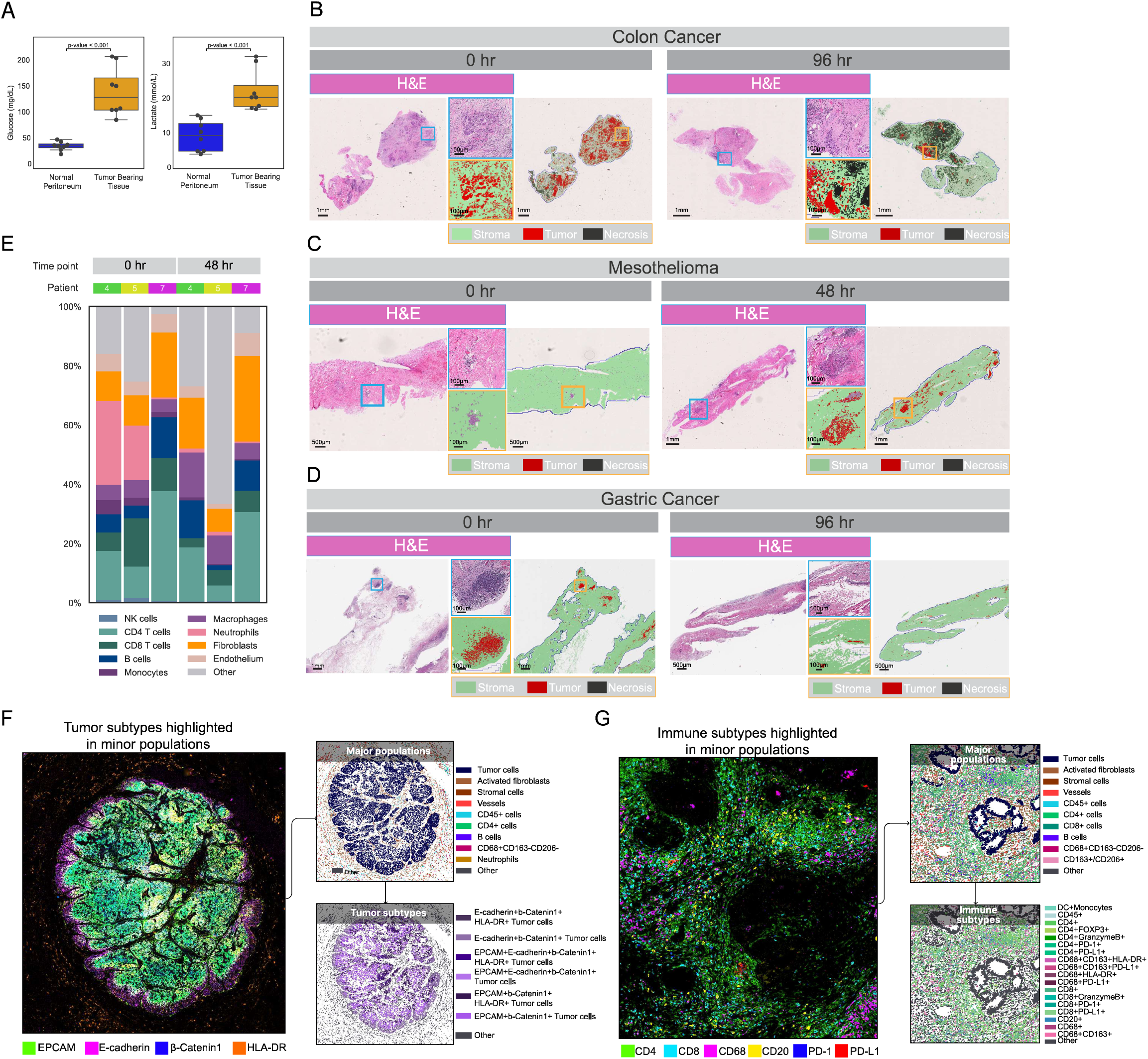
**A:** Boxplots of changes in changes in glucose levels (top) and lactate levels (bottom) in normal peritoneum compared with tumor-bearing peritoneum. Bottom whisker indicates the maximum value or 75th percentile + 1.5 interquartile range (IQR); the top whisker indicates the minimum value or 25th percentile - 1.5 IQR. **B-D:** Pathology analysis on H&E-stained sections of peritoneal metastases from three patients at 0 hrs and at 96 hrs of perfusion. **E:** Bar graphing showing predicted percentages of cell types from transcriptomic-based cell deconvolution in Patient 4, Patient 5, and Patient 7 at 0 hr and 48 hr or 96 hr of perfusion. **F, G:** Representative immunofluorescence and cell typing images to demonstrate cell typing of both major and minor tumor and immune cell populations.

**Figure S4.**
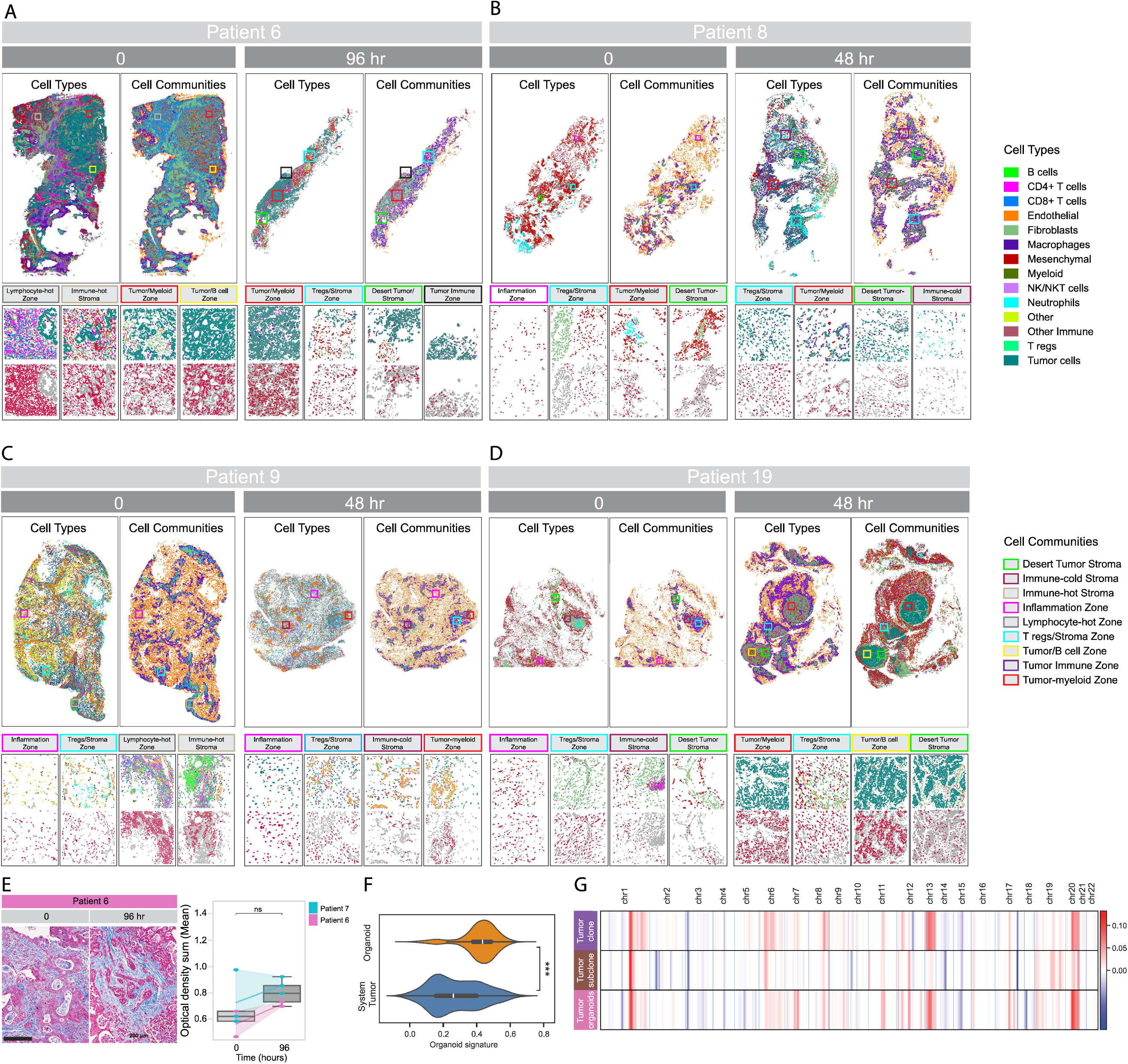
**A-D:** Cell typing and spatial community analysis for tumors of patients 6, 8, 9 and 19 at 0 hr and 48 hr or 96 hr of perfusion. (full tissue-top; tissue regions-bottom). **E:** Representative image of Masson’s Trichrome stain at 0 and 96 hr of perfusion in Patient 6 (left). Boxplots showing cell density at 0 and 96 hrs of perfusion in the system for Patient 6 and Patient 7. In the boxplots, the bottom whisker indicates the maximum value or 75th percentile + 1.5 interquartile range (IQR); the top whisker indicates the minimum value or 25th percentile - 1.5 IQR. **F:** Violin plot of organoid signature applied to organoid (gold) and SMART system perfused tumor (blue) derived from Patient 6. **G:** Heatmap depicting copy number variations on each chromosome in the Patient 6 tumor organoid and Patient 6 SMART system tumors, showing a major clone and subclone in the perfused tumor.

**Figure S5.**
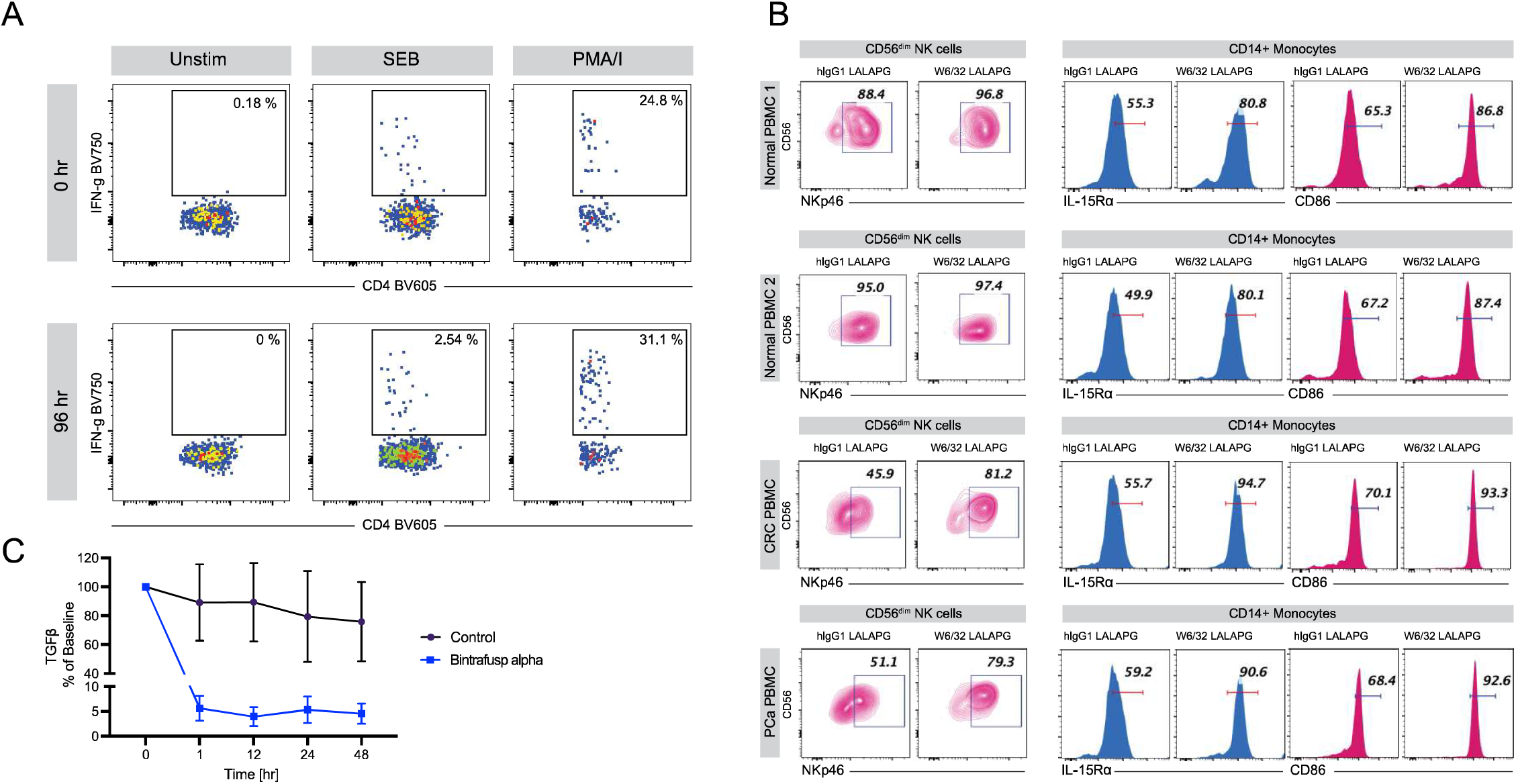
**A:** CD4^+^ T cell responsiveness was assessed at baseline (0 hr, top) and following perfusion (96 hr, bottom) with either SEB, PMA/I, or were unstimulated for 3 hr in the presence of Brefeldin A and were evaluated for IFNγ production. **B:** Flow cytometric analysis of the expression of NKp46 on NK cells, IL-15Ra and CD86 expression on CD14^+^ monocytes in the PBMCs isolated from two healthy donors (normal PBMC 1 and normal PBMC 2) and from two patient PBMCs (colorectal cancer PBMC and pancreatic cancer PBMC) following incubation with control hIgG1LALAPG and W6/32LALAPG monoclonal antibodies for 96 hr. **C:** TGF-β changes in plasma perfusate (% of Ohr) with bintrafusp alpha or control treatment.

