## Supplemental Tables for "A simple circuit to sustain intact tumor microenvironments for complex drug interrogations"

**Supplemental Table S1.**

| Patient Number | Primary Malignancy vs Normal | Tissue Source | Duration of SMART run (hours) | Number of Platforms | Drug Treatment | Key treatment analyzed for paper | Processing | Figure | Treatment |
| --- | --- | --- | --- | --- | --- | --- | --- | --- | --- |
| 1 | Normal | Peritoneum | 48 | 8 | N | SMART system performance | Perfusate & FFPE (pathology, Masson trichrome trial) | N/A | N/A |
| 2 | Normal | Peritoneum | 48 | 8 | N | SMART system performance | Perfusate & FFPE (pathology, Masson trichrome trial) | N/A | N/A |
| 3 | Normal | Peritoneum | 72 | 13 | N | SMART system performance | Perfusate & FFPE (pathology, Masson trichrome trial) | N/A | N/A |
| 4 | CRC | Peritoneal Tumor Deposits | 96 | 4 | N | Bulk transcriptome 0/96hr | RNA sequencing-Snap frozen | 2(A/B), 4(A), 5(M), S3(A) | N/A |
| 5 | Mesothelioma | Peritoneal Tumor Deposits | 96 | 4 | Y | Bulk transcriptome 0/96hr | RNA sequencing-Snap frozen | 2(A/B),4(A), S3(B) | N/A |
| 6 | CRC | Peritoneum and Liver Core Biopsies | 96 | 4 | N | Bulk transcriptome-spatial proteomics-WES-0/96 hr, Single cell seq (SMART vs organoid) | Bulk RNA&CODEX- FFPE, Tissue/organoid dissociation? WES | 2(A/B/E), 3(B/C/D/E/F/G), S5(A/B) | N/A |
| 7 | Gastric Cancer | Peritoneal Tumor Deposits | 96 | 6 | N | Spatial transcriptomics/proteomics 0/96hr | Bulk RNA-FFPE, CODEX & Phospho stains-FFPE | 2(A/B), 4(A/B/C/D), S3(C), S5(A) | N/A |
| 8 | Appendiceal Cancer | Peritoneal Tumor Deposits | 96 | 4 | Y | Bulk transcriptome-spatial proteomics, Dextran 0/96hr (Macrophages), Immune cell populations | CODEX & Bulk RNA- FFPE, Flow Cytometry-Dissociation,Imaging-Fixed wholemount | 2(E), 3(B/C), 4 (H/I), 4 (E-G), S5c | Dextran |
| 9 | Mesothelioma | Peritoneal Tumor Deposits | 48 | 4 | N | Bulk transcriptome/spatial proteomics 0/48hr, Immune cell populations | CODEX&Bulk RNA-FFPE, Dissociation-Flow Cytometry | 2(E), 3(B/C), 4(L/M) | N/A |
| 10 | Cervical Cancer | Tumor Core Biopsies | 48 | 6 | Y | Bulk transcriptome-spatial proteomics/transcriptomics 0/48hr, Cytokines | Bulk RNA&CODEX-FFPE, cytokines-perfusate | 2(B), 5(F/G/H/I/J/K/L/M) | M7824 |
| 11 | Anal Cancer | Liver Core Biopsies | 48 | 4 | Y | Bulk transcriptome-spatial proteomics-WES-0/48 hr, Cytokines | Bulk RNA&CODEX-FFPE, WES, cytokines-perfusate | 2(A/B/C/D), 3(A/B/C), 5(F/G/H/I/J/K/L/M) | M7824 |
| 12 | Mesothelioma | Peritoneal Tumor Deposits | 48 | 16 | Y | Bulk transcriptome-spatial transcriptomics/proteomics-WES-0/48 hr, Cytokines | Bulk RNA&CODEX-FFPE, WES, cytokines-perfusate | 2(A) | M7824 |
| 13 | Gastric Cancer | Peritoneal Tumor Deposits | 48 | 8 | Y | Bulk transcriptome-spatial transcriptomics/proteomics-WES-0/48 hr | Bulk RNA&CODEX-FFPE, WES | 2(A) | N/A |
| 14 | CRC | Peritoneal Tumor Deposits | 48 | 12 | Y | Bulk transcriptome-0/48 hr, Cytokines | Bulk RNA-FFPE, cytokines-perfusate | 2(A/B), 5(M) | M7824 |
| 15 | Mesothelioma | Peritoneal Tumor Deposits | 48 | 12 | Y | Bulk transcriptome-0/48 hr, Cytokines | Bulk RNA-FFPE, cytokines-perfusate | 2(A/B), 5(M) | M7824 |
| 16 | Pancreatic Cancer | Peritoneal Tumor Deposits | 48 | 10 | Y | Bulk transcriptome-0/48 hr, Cytokines | Bulk RNA-FFPE, cytokines-perfusate | 2(A/B), 5(M) | M7824 |
| 17 | Gastric Cancer | Peritoneal Tumor Deposits | 48 | 8 | Y | Bulk transcriptome-0/48 hr, Cytokines | Bulk RNA-FFPE, cytokines-perfusate | 2(A/B), 5(M) | M7824 |
| 18 | Normal | Peritoneum | 96 | 2 | N | SMART system performance (Antibodies) | Perfusate & FFPE (pathologist) |  | N/A |
| 19 | Ovarian Cancer | Peritoneal Tumor Deposits | 48 | 8 | N | SMART system performance (proteomics) | CODEX & Phospho stains-FFPE | 3(C), 4(B/C/D), 5(J/K) | N/A |
| 20 | ACC | Liver Core Biopsies | 96 | 2 | Y | SMART system performance (viability) | Perfusate & FFPE (pathologist) | 3(B) | N/A |
| 21 | Normal | Peritoneum | 96 | 4 | N | SMART system performance (viability) | Perfusate & FFPE (pathologist) |  | N/A |
| 22 | Recurrent Cholangiocarcinoma | Tumor Compresstome Slices | 96 | 12 | Y | SMART system performance (treatment trial) | Perfusate & FFPE (pathologist) |  | N/A |
| 23 | Normal | Peritoneum | 96 | 4 | N | SMART system performance (run duration trial) | Perfusate & FFPE (pathologist) |  | N/A |
| 24 | Normal | Peritoneum | 168 | 4 | N | SMART system performance (run duration trial, contamination) | Perfusate & FFPE (pathologist) |  | N/A |
| 25 | Normal | Peritoneum | 96 | 8 | N | SMART system performance (?) | Perfusate & FFPE (pathologist) |  | N/A |
| 26 | Normal | Peritoneum | 96 | 8 | N | SMART system performance (run duration trial, viability) | Perfusate & FFPE (pathologist) |  | N/A |
| 27 | NET | Peritoneal Tumor Deposits | 96 | 6 | N | SMART system performance (histology/tissue source trial) | Perfusate & FFPE (pathologist) |  | N/A |
| 28 | Normal | Peritoneum | 24 | 7 | N | SMART system performance | Perfusate & FFPE (pathologist) |  | N/A |
| 29 | Normal | Peritoneum | 96 | 8 | N | SMART system performance (dextran) | FFPE (pathologist) |  | N/A |
| 30 | CRC | Liver Compresstome Slices | 48 | 5 | N | SMART system performance (drug treatment ) | Perfusate & FFPE (pathologist) | 5 (A/B/C) | Cetuximab |
| 31 | CRC | Liver Compresstome Slices | 48 | 8 | Y | SMART system performance (drug treatment) | Perfusate & FFPE (pathologist) | 3(B), 5 (A/B/C) | Cetuximab |
| 32 | Normal | Peritoneum | 96 | 2 | N | SMART system performance (immune cells) | FFPE (pathologist) |  | N/A |
| 33 | GIST | Tumor Compresstome Slices | 96 | 4 | N | SMART system performance (histology/tissue source trial) | Perfusate & FFPE (pathologist) |  | N/A |
| 34 | CRC | Liver Compresstome Slices | 24 | 4 | N | SMART system performance (histology/duration trial) | Perfusate & FFPE (pathologist) |  | N/A |
| 35 | CRC | Peritoneal Tumor Deposits | 48 | 12 | Y | SMART system performance (drug treatment optimization) | Perfusate & FFPE (pathologist) |  | N/A |
| 36 | GIST | Peritoneal Tumor Deposits | 96 | 4 | N | SMART system performance (histology trial) | Perfusate & FFPE (pathologist) |  | N/A |
| 37 | PNET | Tumor Core Biopsies | 96 | 4 | N | SMART system performance (histology/duration of run trial) | Perfusate & FFPE (pathologist) |  | N/A |
| 38 | Cholangiocarcinoma | Liver Core Biopsies | 96 | 2 | N | SMART system performance (tissue source trial) | Perfusate & FFPE (pathologist) |  | N/A |
| 39 | Ovarian Cancer | Peritoneal Tumor Deposits | 96 | 5 | Y | SMART system performance | FFPE (pathologist) |  | N/A |
| 40 | Normal | Peritoneum | 72 | 6 | N | SMART system performance | FFPE (pathologist) |  | N/A |
| 41 | Gastric | Peritoneal Tumor Deposits | 48 | 6 | Y | SMART system performance (drug treatment optimization) | Perfusate & FFPE (pathologist) |  | N/A |
| 42 | Ovarian Cancer | Peritoneal Tumor Deposits | 144 | 14 | Y | SMART system performance (drug treatment optimization) | Perfusate & FFPE (pathologist) |  | N/A |
| 43 | Normal | Peritoneum | 96 | 5 | N | SMART system performance (run duration trial) | Perfusate & FFPE (pathologist) |  | N/A |
| 44 | Normal | Peritoneum | 192 | 8 | N | SMART system performance | FFPE nuclear preservation donor/autologous perfusate | 1 (I/J), S1(D) | N/A |
| 45 | PDAC | Tumor Compresstome Slices | 48 | 9 | N | SMART system performance | Perfusate & FFPE (pathologist) |  | N/A |
| 46 | Normal | Peritoneum | 72 | 3 | N | SMART system performance (run duration trial) | Perfusate & FFPE (pathologist) |  | N/A |
| 47 | Mesothelioma | Peritoneal Tumor Deposits | 24 | 6 | Y | SMART system performance (dextran) | FFPE (pathologist) |  | N/A |
| 48 | Normal | Peritoneum | 96 | 2 | N | SMART system performance (run length) | Perfusate & FFPE (pathologist) |  | N/A |
| 49 | Normal | Peritoneum | 96 | 4 | N | SMART system performance (dextran) | FFPE (pathologist) |  | N/A |
| 50 | Normal | Peritoneum | 96 | 4 | N | SMART system performance (dextran) | FFPE (pathologist) |  | N/A |
| 51 | Normal | Peritoneum | 96 | 2 | N | SMART system performance | FFPE (pathologist) |  | N/A |
| 52 | Pheochromocytoma | Peritoneal Tumor Deposits | 48 | 2 | N | SMART system performance (histology trial) | Perfusate & FFPE (pathologist) |  | N/A |
| 53 | CRC | Peritoneal Tumor Deposits | 48 | 4 | N | SMART system performance (run duration trial, viability) | Perfusate & FFPE (pathologist) |  | N/A |
| 54 | Rectal Cancer | Liver Core Biopsies | 96 | 10 | Y | SMART system performance (core biopsies) | Perfusate & FFPE (pathologist) |  | N/A |
| 55 | Normal | Peritoneum | 96 | 8 | N | SMART system performance (stir bar) | Perfusate & FFPE (pathologist) |  | N/A |
| 56 | Gastric Cancer | Peritoneal Tumor Deposits | 48 | 8 | Y | SMART system performance (histology trial) | Perfusate & FFPE (pathologist) |  | N/A |
| 57 | Normal | Peritoneum | 96 | 18 | N | SMART system performance (nuclear loss) | Perfusate & FFPE (pathologist) |  | N/A |
| 58 | Mucinous Appendiceal | Peritoneal Tumor Deposits | 96 | 4 | Y | SMART system performance (too mucinous) | Perfusate & FFPE (pathologist) |  | N/A |
| 59 | Granulosa Tumor | Peritoneal Tumor Deposits | 120 | 12 | Y | SMART system performance (treatment trial) | Perfusate & FFPE (pathologist) |  | N/A |
| 60 | CRC | Liver Compresstome Slices | 96 | 4 | N | SMART system performance (slices) | Perfusate & FFPE (pathologist) |  | N/A |
| 61 | Ovarian Cancer | Peritoneal Tumor Deposits | 96 | 6 | N | SMART system performance (trial immune cell population) | Perfusate & FFPE (pathologist) |  | N/A |
| 62 | Mesothelioma | Peritoneal Tumor Deposits | 48 | 7 | N | SMART system performance | Perfusate & FFPE (pathologist) |  | N/A |
| 63 | Pancreatic cancer | Peritoneal Tumor Deposits | 48 | 12 | Y | SMART system performance (treatment trial) | Perfusate & FFPE (pathologist) |  | N/A |
| 64 | CRC | Peritoneal Tumor Deposits | 96 | 8 | N | SMART system performance (stir bar) | Perfusate & FFPE (pathologist) |  | N/A |
| 65 | Mesothelioma | Peritoneal Tumor Deposits | 48 | 4 | Y | SMART system performance (treatment trial) | Flow Cytometry-Dissociation |  | N/A |
| 66 | Normal | Peritoneum | 96 | 7 | N | SMART system performance (nuclear loss) | Perfusate & FFPE (pathologist) |  | N/A |
| 67 | Mesothelioma | Tumor Core Biopsies | 48 | 2 | Y | SMART system performance (treatment trial) | Perfusate & FFPE (pathologist) |  | N/A |
| 68 | Normal | Peritoneum | 96 | 8 | N | SMART system performance (perfusate) | Perfusate & FFPE (pathologist) |  | N/A |
| 69 | ACC | Peritoneal Tumor Deposits | 96 | 4 | N | SMART system performance (histology trial) | Perfusate & FFPE (pathologist) |  | N/A |
| 70 | Normal | Peritoneum | 96 | 2 | N | SMART system performance | Perfusate & FFPE (pathologist) |  | N/A |
| 71 | CRC | Peritoneal Tumor Deposits | 24 | 6 | N | SMART system performance (treatment trial) | Perfusate & FFPE (pathologist) |  | N/A |
| 72 | GIST | Peritoneal Tumor Deposits | 96 | 12 | Y | SMART system performance (treatment trial) | Perfusate & FFPE (pathologist) |  | N/A |
| 73 | Pheochromocytoma | Peritoneal Tumor Deposits | 96 | 4 | N | SMART system performance (histology trial) | Perfusate & FFPE (pathologist) |  | N/A |
| 74 | Normal | Peritoneum | 216 | 5 | N | SMART system performance | Perfusate & FFPE (pathologist) |  | N/A |
| 75 | Normal | Peritoneum | 96 | 8 | N | SMART system performance | Perfusate & FFPE (pathologist) |  | N/A |
| 76 | Normal | Peritoneum | 96 | 4 | N | SMART system performance | Perfusate & FFPE (pathologist) |  | N/A |
| 77 | Normal | Peritoneum | 96 | 3 | N | SMART system performance | Perfusate & FFPE (pathologist) |  | N/A |
| 78 | CRC | Liver Core Biopsies | 96 | 8 | Y | SMART system performance (treatment trial) | Perfusate & FFPE (pathologist) |  | N/A |
| 79 | Normal | Peritoneum | 48 | 8 | N | SMART system performance (perfusate additives) | FFPE (pathologist) |  | N/A |
| 80 | Normal | Peritoneum | 96 | 2 | N | SMART system performance (Antibodies) | Perfusate & FFPE (pathologist) |  | N/A |
| 81 | Gastric Cancer | Peritoneal Tumor Deposits | 96 | 3 | N | SMART system performance (run duration trial, viability) | Perfusate & FFPE (pathologist) |  | N/A |
| 82 | Normal | Peritoneum | 48 | 8 | N | SMART system performance (perfusate additives) | FFPE (pathologist) |  | N/A |
| 83 | Normal | Peritoneum | 96 | 8 | N | SMART system performance | Perfusate & FFPE (pathologist) |  | N/A |
| 84 | Normal | Peritoneum | 48 | 4 | N | SMART system performance (PDMS) | Perfusate & FFPE (pathologist) |  | N/A |
| 85 | Normal | Peritoneum | 96 | 8 | N | SMART system performance (perfusate) | Perfusate & FFPE (pathologist) |  | N/A |
| 86 | CRC | Peritoneal Tumor Deposits | 96 | 6 | N | SMART system performance (histology trial) | Perfusate & FFPE (pathologist) |  | N/A |
| 87 | CRC | Peritoneal Tumor Deposits | 48 | 7 | Y | SMART system performance (treatment trial) | Perfusate & FFPE (pathologist) |  | N/A |
| 88 | CRC | Peritoneal Tumor Deposits | 96 | 1 | N | SMART system performance (run duration trial, viability) | Perfusate & FFPE (pathologist) |  | N/A |
| 89 | Normal | Peritoneum | 120 | 5 | N | SMART system performance (run duration trial, viability) | Perfusate & FFPE (pathologist) |  | N/A |
| 90 | Normal | Peritoneum | 96 | 5 | N | SMART system performance (run duration trial, viability) | Perfusate & FFPE (pathologist) |  | N/A |
| 91 | Normal | Peritoneum | 48 | 2 | N | SMART system performance (nuclear preservation) | Perfusate & FFPE (pathologist) |  | N/A |
| 92 | CRC | Liver Compresstome Slices | 96 | 6 | N | SMART system performance (slices) | Perfusate & FFPE (pathologist) |  | N/A |
| 93 | Normal | Peritoneum | 96 | 10 | N | SMART system performance (perfusate) | Perfusate & FFPE (pathologist) |  | N/A |
| 94 | Normal | Peritoneum | 96 | 2 | N | SMART system performance (run duration trial) | Perfusate & FFPE (pathologist) |  | N/A |
| 95 | Gastric Cancer | Peritoneal Tumor Deposits | 48 | 2 | Y | SMART system performance (treatment trial) | Perfusate & FFPE (pathologist) |  | N/A |
| 96 | NET | Liver Core Biopsies | 96 | 4 | N | SMART system performance (histology trial) | Perfusate & FFPE (pathologist) |  | N/A |
| 97 | Normal | Peritoneum | 96 | 8 | N | SMART system performance (perfusate) | Perfusate & FFPE (pathologist) |  | N/A |
| 98 | Normal | Peritoneum | 168 | 14 | N | SMART system performance (run duration trial) | Perfusate & FFPE (pathologist) |  | N/A |
| 99 | Normal | Peritoneum | 120 | 5 | N | SMART system performance (run duration trial) | Perfusate & FFPE (pathologist) |  | N/A |
| 100 | Gastric Cancer | Peritoneal Tumor Deposits | 96 | 14 | N | Wholemount Imaging Trial | FFPE (pathologist) |  | N/A |
| 101 | Gastric Cancer | Peritoneal Tumor Deposits | 192 | 8 | N | SMART system performance (run duration trial) | Perfusate & FFPE (pathologist) |  | N/A |
| 102 | ACC | Tumor Core Biopsies | 48 | 3 | N | SMART system performance (histology trial) | Perfusate & FFPE (pathologist) |  | N/A |
| 103 | CRC | Liver Compresstome Slices | 96 | 4 | Y | SMART system performance (treatment trial) | Perfusate & FFPE (pathologist) |  | N/A |
| 104 | RCC | Tumor Core Biopsies | 96 | 6 | Y | SMART system performance (histology&treatment trial) | Perfusate & FFPE (pathologist) |  | N/A |
| 105 | Appendiceal adenocarcinoma | Peritoneal Tumor Deposits | 48 | 2 | Y | SMART system performance (treatment trial) | Perfusate & FFPE (pathologist) |  | N/A |
| 106 | PNET | Liver Compresstome Slices | 96 | 8 | N | SMART system performance (histology&run duration trial) | Perfusate & FFPE (pathologist) |  | N/A |
| 107 | Gastric Cancer | Peritoneal Tumor Deposits | 120 | 20 | N | SMART system performance (run duration trial) | Perfusate & FFPE (pathologist) |  | N/A |
| 108 | Normal | Peritoneum | 96 | 2 | N | SMART system performance (run duration trial) | Perfusate & FFPE (pathologist) |  | N/A |
| 109 | Cholangiocarcinoma | Liver Core Biopsies | 96 | 6 | Y | SMART system performance (histology trial) | Perfusate & FFPE (pathologist) |  | N/A |
| 110 | CRC | Liver Core Biopsies | 96 | 16 | Y | SMART system performance (treatment trial) | Perfusate & FFPE (pathologist) |  | N/A |
| 111 | Normal | Peritoneum | 96 | 2 | N | SMART system performance (perfusate) | Perfusate & FFPE (pathologist) |  | N/A |
| 112 | Mesothelioma | Peritoneal Tumor Deposits | 48 | 9 | Y | SMART system performance (treatment trial) | Perfusate & FFPE (pathologist) |  | N/A |
| 113 | Normal | Peritoneum | 96 | 7 | N | SMART system performance (run duration trial) | Perfusate & FFPE (pathologist) |  | N/A |
| 114 | Mesothelioma | Peritoneal Tumor Deposits | 96 | 8 | Y | SMART system performance (treatment trial) | Perfusate & FFPE (pathologist) |  | N/A |
| 115 | Normal | Peritoneum | 96 | 4 | N | Dextran, Macrophages | Fixed wholemount Imaging | 1 (K/L) | Dextran |
| 116 | Appendiceal Cancer | Peritoneal Tumor Deposits | 96 | 4 | N | SMART system performance (histology trial) | Perfusate & FFPE (pathologist) |  | N/A |
| 117 | Mesothelioma | Peritoneal Tumor Deposits | 48 | 4 | Y | SMART system performance (treatment trial) | Perfusate & FFPE (pathologist) |  | N/A |
| 118 | Mesothelioma | Peritoneal Tumor Deposits | 48 | 6 | Y | SMART system performance (treatment trial) | Perfusate & FFPE (pathologist) |  | N/A |
| 119 | Normal | Peritoneum | 120 | 8 | N | SMART system performance (duration/platform trial) | Perfusate & FFPE (pathologist) |  | N/A |
| 120 | Mesothelioma | Peritoneal Tumor Deposits | 48 | 3 | Y | SMART system performance (treatment trial) | Perfusate & FFPE (pathologist) |  | N/A |
| 121 | CRC | Peritoneal Tumor Deposits | 48 | 4 | Y | SMART system performance (treatment trial) | Perfusate & FFPE (pathologist) |  | N/A |
| 122 | CRC | Liver Core Biopsies | 48 | 8 | Y | SMART system performance (treatment trial) | Perfusate & FFPE (pathologist) |  | N/A |
| 123 | Normal | Peritoneum | 120 | 5 | N | SMART system performance (run duration trial) | Perfusate & FFPE (pathologist) |  | N/A |
| 124 | Mesothelioma | Peritoneal Tumor Deposits | 72 | 14 | Y | SMART system performance (treatment trial) | Perfusate & FFPE (pathologist) |  | N/A |
| 125 | CRC | Peritoneal Tumor Deposits | 96 | 8 | Y | SMART system performance (treatment trial) | Perfusate & FFPE (pathologist) |  | N/A |
| 126 | Mesothelioma | Peritoneal Tumor Deposits | 48 | 10 | Y | SMART system performance (treatment trial) | Perfusate & FFPE (pathologist) |  | N/A |
| 127 | Gastric | Peritoneal Tumor Deposits | 72 | 2 | Y |  | Phospho stains-FFPE | 4 (J/K) | Bintrafusp alfa |
| 128 | CRC | Liver Core Biopsies | 48 | 4 | Y | W6/32, NK cells | Flow Cytometry-Dissociation | 5(D/E), S5(D) | W6/32 |
| 129 | Pancreatic cancer | Liver Core Biopsies | 48 | 4 | Y | W6/32, NK cells | Flow Cytometry-Dissociation | 5(D/E), S5(D) | W6/32 |

Supplemental Table 1: Patient samples

**Supplemental Table S2.**

| **3D-printing of System Components** | | | |
| --- | --- | --- | --- |
| Company/Vendor | Part | Cat Number | purpose |
| FormLabs | BioMed Clear Resin (Form 3) | RS-F2-BMCL-01 | resin used for platforms and culture dish |
| Formlabs | Form 3B+ | Form 3B+ | printer used for platform + incubation plate |
| Formlabs | Form Cure | FH-CU-01 | post processing for platform + incubation plate |
| Formlabs | Form Wash | FH-WA-01 | post processing for platform + incubation plate |
| FormLabs | PreForm | PreForm | Software to prepare models for Formlab printers |
| **Smart System Set up** | | | |
| Company/Vendor | Part | Cat Number | purpose |
| McMaster-Carr | Polycarbonate Plugs, Barbed for 1/8" Tube ID | 51525K273 | One used per SMART system set up |
| VWR | Masterflex® Ismatec® Pump Tubing, 3-Stop, Puri-Clear™ LL, 2.79 mm ID; 12/PK | MFLX95625-48 | Tubing to circulate SMART system perfusate |
| Cole Palmer | Ismatec MS/CA Click-N-Go Cassette Cartridge, 3-Stop, POM-C | SK-95625-48 | Cassette which clicks into the pump to allow circulation of perfusate |
| Masterflex | Ismatec Reglo Digital Pump with MasterflexLive™, 4-Channel, 8-Roller; 115/230 VAC | EW-78016-98 | Pump that actually circulates the perfusate |
| PermSelect | PDMSXA-100: PermSelect® 100 centimeter sq. membrane module with barb fitting | HV-78018-22 | Oxygenator for SMART system |
| McKesson | Stopcock Discofix® 4 Way | 165802 | Two utilized for each SMART system |
| Masterflex | Masterflex® Ismatec® Microbore Pump Tubing, Peroxide-Cured Silicone, Avantor® | MFLX07625-48 | Tubing utilized to deliver gas mix into oxygenator from gasmixer |
| Medex Supply | Clave neutral connector | ICU-12568 | port to take samples for labs |
| Medfusion | syringe pumps | Medfusion 3500, G600-736 | free water delivery |
| MCQ Instruments | Gasmixer | GB100plus | generate CO2/O2 gas mix |
| Robert's Oxygen | Carbon Dioxide Tank with regulator | R25, CUF1YT1 | generate CO2/O2 gas mix |
| Robert's Oxygen | Oxygen Tank with regulator | R5, CUF1YT1 | generate CO2/O2 gas mix |
| BD | BD Disposable Syringes with Luer-Lok™ Tips, 1mL | 309628 | take labs |
| BD | 10 mL BD Luer-Lok™ Syringe | 302995 | mount on syringe pumps |
| 3M | 3M Steri drape; large towel drape | 3M-1010-BOX | incubator lining |
| Grayline Medical | Vyaire Humidifiers Prefilled with Sterile Water | 2620 | humidify gas mix |
| **Tissue Mounting/Preparation** | | | |
| Company/Vendor | Part | Cat Number | purpose |
| Ethicon | Silk sutures | A305.O35 | affix tissue to platform |
| Spring USA | Warming Tray | ST-1220 | keep media and tissue warm |
| Falcon | Dish 15 cm | 353025 | dish used during mounting |
| McKesson | Container, Specimen | 870203 | transfer of tissue from perating table to prep table |
| Gibco | DMEM | 11966025 | harvest medium |
| Gibco | Antibiotic-Antimycotic | 15240096 | supplement harvest medium |
| Precisionary | Vibratome | VF-310-0Z | preparation of tissue sections |
| **Perfusate Preparation** | | | |
| Company/Vendor | Part | Cat Number | purpose |
| Millipore Sigma | 50cc conical tube | CLS430290-500EA | Utilized to isolate plasma from blood |
| Gibco | Antibiotic-Antimycotic | 15240096 | supplement perfusate |
| Fresenius Kabi | Heparin Sodium Injection, USP, 50,000 USP units per 10 mL (5,000 USP units per mL) | 4710 | Utilized to isolate plasma from blood |
| Novo Nordisk Pharmaceutical | Novolin® R Regular Human Insulin (rDNA Origin) 100 U / mL Injection 10 mL | 169183311 | supplement perfusate |
| B. Braun | 10% Dextrose Injection, USP | L5202 | supplement perfusate |
| **SMART System Monitoring** | | | |
| Company/Vendor | Part | Cat Number | purpose |
| Nova Biomedical | Stat Profile Prime Plus® CCS, Analyzer | 59423 | Point-of-Care Meter |
| Nova Biomedical | Microsensor Cartridge | 58642 | supplies for Point-of-Care Meter |
| Vet One | Sodium Bicarbonate 8.4% Injection | 510186 | supplement perfusate |
| B. Braun | Sterile Water for Irrigation USP, 1000 mL | R5000-01 | supplement perfusate |
| B. Braun | 10% Dextrose Injection, USP | L5202 | supplement perfusate |

Supplemental Table 2: Experimental materials

**Supplementary Table S3.**

| **CODEX** | | | | | | |
| --- | --- | --- | --- | --- | --- | --- |
| Company | Cat No | Target | Clone | Reporter Dye | Dilution | Comments |
| Akoya | 4450036 | b-Catenin1 | 12F7 | Atto 550 | 1:200 |  |
| Akoya | 4250098 | Bcl-2 | EPR17509 | Alexa Fluor 647 | 1:200 |  |
| Akoya | 4450040 | Beta-actin | W16197A | Alexa Fluor 750 | 1:200 |  |
| Akoya | 4550119 | CD3e | EP449E | Alexa Fluor 647 | 1:200 |  |
| Akoya | 4550112 | CD4 | EPR6855 | Alexa Fluor 647 | 1:200 |  |
| Akoya | 4250012 | CD8 | C8/144B | Atto 550 | 1:200 |  |
| Akoya | 4450018 | CD20 | L26 | Alexa Fluor 750 | 1:200 |  |
| Akoya | 4450027 | CD21 | EP3093 | Atto 550 | 1:200 |  |
| Akoya | 4450017 | CD31 | EP3095 | Alexa Fluor 750 | 1:200 |  |
| Akoya | 4250057 | CD34 | QBEND10 | Atto 550 | 1:200 |  |
| Akoya | 4250080 | CD38 | E7Z8C | Atto 550 | 1:200 |  |
| Akoya | 4250076 | CD39 | EPR20627 | Atto 550 | 1:200 |  |
| Akoya | 4450041 | CD44 | 156-3C11 | Atto 550 | 1:200 |  |
| Akoya | 4450042 | CD45 | D9M81 | Cy5 | 1:200 |  |
| Akoya | 4250023 | CD45RO | UCHL1 | Atto 550 | 1:200 |  |
| Akoya | 4550001 | CD66 | ASL-32 | Alexa Fluor 647 | 1:200 |  |
| Akoya | 4550113 | CD68 | KP1 | Alexa Fluor 647 | 1:200 |  |
| Akoya | 4450078 | CD79a | D1X5C | Alexa Fluor 750 | 1:200 |  |
| Akoya | 4550098 | CD107a | H4A3 | Alexa Fluor 647 | 1:200 |  |
| Akoya | 4250097 | CD141 | E7Y9P | Atto 550 | 1:200 |  |
| Akoya | 4250079 | CD163 | EPR19518 | Atto 550 | 1:200 |  |
| Akoya | 4450122 | Collagen IV | EPR209660 | Alexa Fluor 647 | 1:200 |  |
| Akoya | 4250021 | E-cadherin | 4A2C7 | Atto 550 | 1:200 |  |
| Akoya | 4550088 | EpCAM | D9S3P | Alexa Fluor 647 | 1:200 |  |
| Akoya | 4550071 | FOXP3 | 236A/E7 | Alexa Fluor 647 | 1:200 |  |
| Akoya | 4250055 | Granzyme B | D6E9W | Atto 550 | 1:200 |  |
| Akoya | 4450046 | HLA-A | EP1395Y | Alexa Fluor 750 | 1:200 |  |
| Akoya | 4550118 | HLA-DR | EPR3692 | Alexa Fluor 647 | 1:200 |  |
| Akoya | 4250065 | HLA-E | MEM- E/02 | Atto 550 | 1:200 |  |
| Akoya | 4550117 | ICOS | D1K2T | Alexa Fluor 647 | 1:200 |  |
| Akoya | 4250062 | IFNG | EPR21704 | Atto 550 | 1:200 |  |
| Akoya | 4250019 | Ki67 | B56 | Atto 550 | 1:200 |  |
| Akoya | 4250083 | MPO | E1E7I | Atto 550 | 1:200 |  |
| Akoya | 4450020 | Pan-Cytokeratin | AE-1/AE-3 | Alexa Fluor 750 | 1:200 |  |
| Akoya | 4550124 | PCNA | PC10 | Alexa Fluor 647 | 1:200 |  |
| Akoya | 4550038 | PD-1 | D4W2J | Alexa Fluor 647 | 1:200 |  |
| Akoya | 4550072 | PD-L1 | 73-10 | Alexa Fluor 647 | 1:120 |  |
| Akoya | 4450049 | aSMA | 1A4 | Alexa Fluor 750 | 1:200 |  |
| Akoya | 4550081 | TP63 | W15093A | Alexa Fluor 647 | 1:200 |  |
| Akoya | 4450050 | Vimentin | 091D3 | Alexa Fluor 750 | 1:300 |  |
| Akoya | 4550069 | HIF1-alpha | EP1215Y | Alexa Fluor 647 | 1:200 |  |
| BD Pharmingen | 555486 | CD45RA | HI100 | Alexa Fluor 647 | 1:100 | custom conjugated |
| BD Pharmingen | 567710 | CD15 | HI98 | Alexa Fluor 647 | 1:150 | custom conjugated |
| CD16 | 72204 | CD16 | D1N9L | Atto 550 | 1:150 | custom conjugated |
| Ultivue | | | | | | |
| Company | Cat No | Target | Clone | Reporter Dye |  | Comment |
| Vizgen | ULT00136 | CD20 | L26 | FITC | 1:100 |  |
| Vizgen | ULT00145 | GranzymeB | EPR8260 | TRITC | 1:100 |  |
| Vizgen | ULT00202 | CD56 | 3H15L12 | Cy5 | 1:100 |  |
| Vizgen | ULT00164 | CD163 | EPR19518 | Cy7 | 1:100 |  |
| Vizgen | ULT00139 | CD8 | C8/144B | FITC | 1:100 |  |
| Vizgen | ULT00138 | PD-1 | CAL20 | TRITC | 1:100 |  |
| Vizgen | ULT00142 | PD-L1 | 73-10 | Cy5 | 1:100 |  |
| Vizgen | ULT00137 | CD68 | KP-1 | Cy7 | 1:100 |  |
| Vizgen | ULT00140 | CD3 | BC33 | FITC | 1:100 |  |
| Vizgen | ULT00143 | CD4 | SP35 | TRITC | 1:100 |  |
| Vizgen | ULT00292 | FoxP3 | 236A/E7 | Cy5 | 1:100 |  |
| Vizgen | ULT00135 | CK/SOX10 | AE1/AE3 & BC34 | Cy7 | 1:100 |  |
| **GeoMx** | | | | | | |
| Company | Cat No | Target | Clone | Reporter Dye |  | Comment |
| BioRad | MCA5781GA | SMA | 1A4 | Alexa Fluor 647 | 1:200 | custom conjugated, Abcam (Ab269823) |
| Nanostring | 121300310 | panCK |  | Alexa Fluor 532 | 1:40 | GeoMx Solid Tumor Morphology Kit |
| Nanostring | 121300310 | CD45 | 2B11+PD7/26 | Alexa Fluor 594 | 1:40 | GeoMx Solid Tumor Morphology Kit |
| Nanostring | 121300303 | Syto 13 (nuclear stain) | N/A |  | 1:10 |  |
| **Flow Cytometry II (W6/32)** | | | | | | |
| Company | Cat No | Target | Clone | Reporter Dye |  | Comment |
| BioLegend | 362554 | CD56 | 5.1H11 | APC/Fire |  |  |
| BioLegend | 302040 | CD16 | 3G8 | BV605 |  |  |
| BioLegend | 331927 | CD335(NKp46) | 9E2 | BV650 |  |  |
| BD Biosciences | 560957 | CD86 | 2331 | PE |  |  |
| BD Biosciences | 612750 | CD3 | UCHT1 | BUV737 |  |  |
| BD Biosciences | 563743 | CD14 | MΦP9 | BV421 |  |  |
| R&D systems | FAB1471T | IL-15R alpha | N/A | Alexa Fluor 594 |  |  |
| Thermo Fisher | 53-0199-42 | CD19 | HIB19 | Alexa Fluor 488 |  |  |
| **Flow Cytometry I (48/96 hr IME)** | | | | | | |
| Company | Cat No | Target | Clone | Reporter Dye |  | Comment |
| BD Biosciences | 568335 | HLA-DR | G46-6 | BUV805 |  | Extracellular |
| BD Biosciences | 741603 | CD14 | M5E2 | BUV661 |  | Extracellular |
| BD Biosciences | 752378 | CD15 | 7C3.rMAb | BV711 |  | Extracellular |
| BD Biosciences | 744437 | CD11c | S-HCL-3 | BV650 |  | Extracellular |
| BD Biosciences | 566357 | IFNg | B27 | BV750 |  | Intracellular |
| BioLegend | 362508 | CD56 | 5.1H11 | PerCP-Cy5.5 |  | Extracellular |
| BioLegend | 302025 | CD16 | 3G8 | ALX700 |  | Extracellular |
| BioLegend | 302230 | CD19 | HIB19 | PerCP-Cy5.5 |  | Extracellular |
| BioLegend | 305110 | CD66b | G10F5 | ALX647 |  | Extracellular |
| BioLegend | 300306 | CD3 | HIT3a | FITC |  | Intracellular |
| BioLegend | 317438 | CD4 | OKT4 | BV605 |  | Intracellular |
| BioLegend | 301046 | CD8 | RPA-T8 | BV785 |  | Intracellular |
| BioLegend | 320116 | FoxP3 | 206D | Pacific Blue |  | Intracellular |
| R&D Systems | FAB1430T | CD45 | 2D1 | ALX594 |  | Extracellular |
| ThermoFisher | L34962 | Amine-reactive dye | N/A | LIVE/DEAD™ Fixable Blue Dead Cell Stain Kit, for UV excitation |  | Extracellular |
| **MCS primary Antibodies (IF, IHC)** | | | | | | |
| Company | Cat No | Target | Clone | Reporter Dye |  | Comment |
| Abcam | ab133616 | CD4 | EPR6855 | N/A | 1:250 |  |
| Abcam | ab228462 | PD-L1 | SP142 | N/A | 1:100 |  |
| Abcam | ab233396 | CD69 | EPR21814 | N/A | 1:250 |  |
| Abcam | ab101500 | CD8 alpha | SP16 | N/A | 1:100 |  |
| Biolegend | 307620 | HLA-DR | L243 | Alexa Fluor 488 | 1:130 |  |
| BioRad | MCA1477 | CD3 | CD3-12 | N/A | 1:100 |  |
| Cell Signaling | 9145 | pStat3 (Tyr705) | D3A7 | N/A | 1:50 |  |
| Cell Signaling | 4857 | pS6 Ribosomal Protein (Ser235/236) | 91B2 | N/A | 1:150 |  |
| Cell Signaling | 3787 | pAkt (Ser473) | 736E11 | N/A | 1:50 |  |
| Cell Signaling | 4376 | p44/42 MAPK (Erk1/2) (Thr202/Tyr204) | 20G11 | N/A | 1:200 |  |
| Cell Signaling | 9661 | Cleaved Caspase-3 (Asp175) | N/A | N/A | 1:100 |  |
| Cell Signaling | 37259 | CD44 | E7K2Y | N/A | 1:400 |  |
| Cell Signaling | 3195 | E-Cadherin | 24E10 | N/A | 1:400 |  |
| Cell Signaling | 13917 | CD45 | D9M8I | N/A | 1:50 |  |
| Cell Signaling | 9027 | Ki-67 | D2H10 | N/A | 1:100 |  |
| Cell Signaling | 99746 | NCAM1 (CD56) | E7X9M | N/A | 1:50 |  |
| DAKO/Agilent | M0876 | CD68 | PG-M1 | N/A | 1:50 |  |

Supplemental Table 3: Antibodies
